# Degeneracy and stability in neural circuits of dopamine and serotonin neuromodulators: A theoretical consideration

**DOI:** 10.1101/2020.09.25.313999

**Authors:** Chandan K. Behera, Alok Joshi, Da-Hui Wang, Trevor Sharp, KongFatt Wong-Lin

## Abstract

Degenerate neural circuits perform the same function despite being structurally different. However, it is unclear whether neural circuits with interacting neuromodulator sources can themselves be degenerate while maintaining the same neuromodulatory function. Here, we address this by computationally modelling the neural circuits of neuromodulators serotonin and dopamine, local glutamatergic and GABAergic interneurons, and their possible interactions, under reward/punishment-based conditioning tasks. The neural modelling is constrained by relevant experimental studies of the VTA or DRN system using e.g. electrophysiology, optogenetics, and voltammetry. We first show that a single parsimonious, sparsely connected neural circuit model can recapitulate several separate experimental findings that indicated diverse, heterogeneous, distributed and mixed DRN-VTA neuronal signalling in reward and punishment tasks. The inability for this model to recapitulate all observed neuronal signalling suggests potentially multiple circuits acting in parallel. Then using computational simulations and dynamical systems analysis, we demonstrate that several different stable circuit architectures can produce the same observed network activity profile, hence demonstrating degeneracy. Due to the extensive D2-mediated connections in the investigated circuits, we simulate D2 receptor agonist by increasing the connection strengths emanating from the VTA DA neurons. We found that the simulated D2 agonist can distinguish among sub-groups of the degenerate neural circuits based on substantial deviations in specific neural populations’ activities in reward and punishment conditions. This forms a testable model prediction using pharmacological means. Overall, this theoretical work suggests the plausibility of degeneracy within neuromodulator circuitry and has important implication for the stable and robust maintenance of neuromodulatory functions.

## 1 Introduction

The nervous system can be modulated by endogenous chemical messengers, called neuromodulators (Levitan, 1987). Neurons or synapses, and hence neural circuits, that succumb to neuromodulation can change circuit configuration and function (Marder, 2012; Marder et al., 2014a). It is also known that neural circuits can be degenerate, that is, circuits can comprise different elements and/or structure while performing the same function or yielding the same output (Cropper et al., 2016). This can allow robust maintenance of functions and behavior in the face of changes in the underlying structure (Edelman and Gally, 2001; Whitacre, 2010). It should be noted that the degeneracy defined here should be distinguished from other definitions e.g. having different states with the same energy level in quantum physics or decline in health condition (Schroeter and Griffiths, 2018; Hocking et al., 2019). Although it has been shown that neuromodulators can selectively regulate degenerate neural circuits (Marder et al., 2014b; Cropper et al., 2016), it is unclear whether neural circuits with neuromodulator-containing neurons can themselves be degenerate, which could in turn provide stable widespread neuromodulator influences on targeted neural circuits (Fig. 1A).

**Figure 1.**
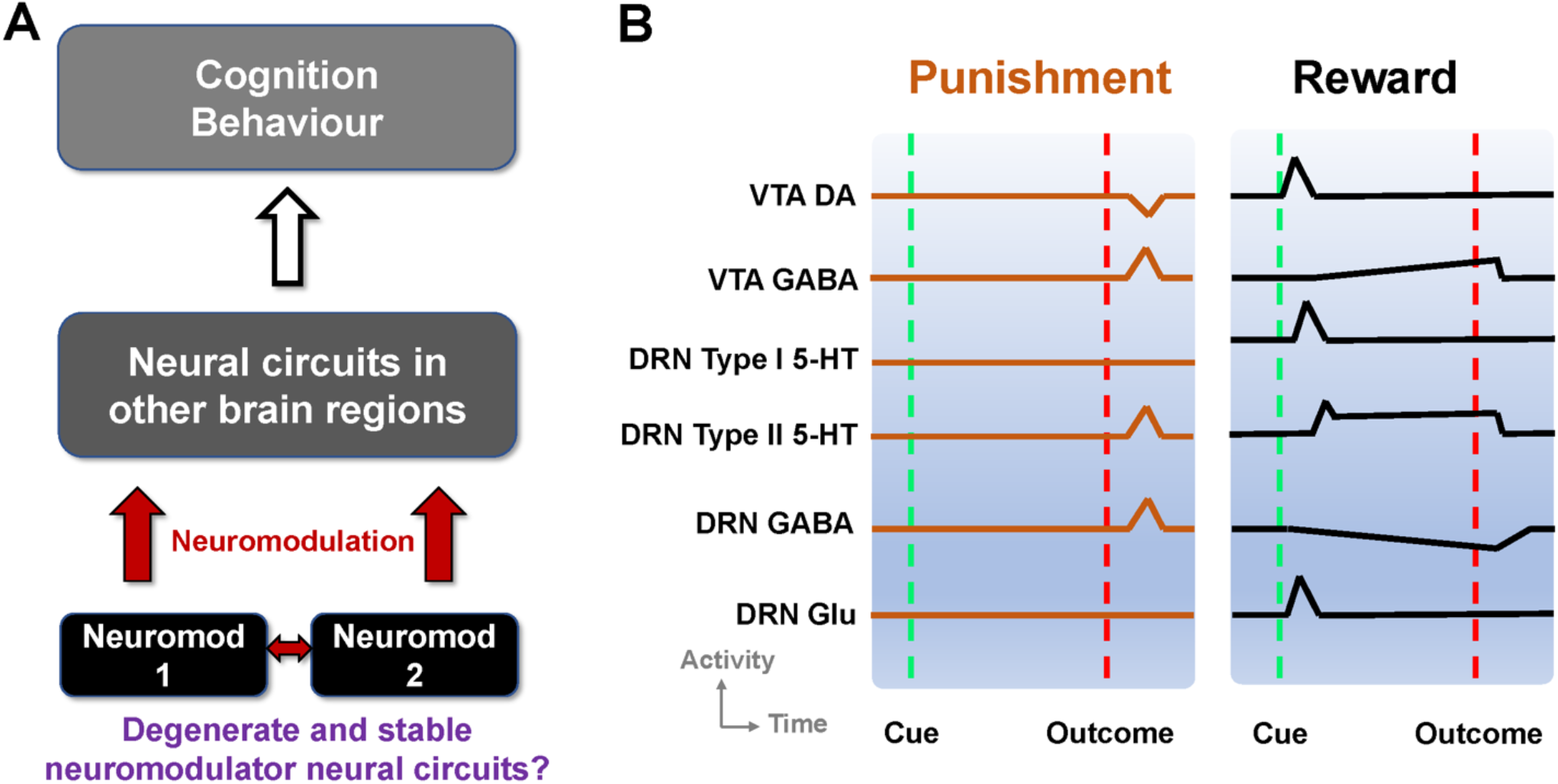
Degenerate neuromodulator circuits constrained by observed neuronal signalling. **A)** Multiple neuromodulators that influence neural circuits, cognition and behaviour, may be embedded within degenerate neural circuits. Neuromod: Specific neuromodulator type. **B)** Schematic of DRN and VTA activity profiles in reward and punishment tasks. Activities (firing rates) aligned to timing of unexpected punishment outcome (left, vertical red dashed lines) and learned reward-predictive cue (right, vertical green dashed lines) and reward outcome (right, vertical red dashed lines). Top-to-bottom: VTA DA neural activity exhibits phasic excitation (inhibition) at reward-predictive cue (punishment) outcome (e.g. (Cohen and Uchida, 2012; Tan et al., 2012)]). VTA GABAergic neural activity shows phasic excitation upon punishment (e.g. (Tan et al., 2012; Eshel et al., 2015)), while exhibiting post-cue tonic activity which is not modulated by the presence/absence of actual outcome (e.g. (Cohen and Uchida, 2012)). DRN Type-I 5-HT neurons shows phasic activation by reward-predicting cue (right) but not punishment (left). DRN Type-II 5-HT neurons signal punishment outcome (left) and sustained activity towards expected reward outcome (right) (e.g. (Cohen et al., 2015b)). DRN GABAergic neurons have phasic activation upon punishment but have tonic inhibition during waiting and reward delivery (e.g. (Li et al., 2016)). DRN glutamatergic neurons deduced to be excited by reward-predicting cue, in line with VTA DA neural activation (McDevitt et al., 2014), and assumed not to respond to punishment outcome. Baseline activity for Type-I 5-HT DRN neurons is higher in reward than punishment tasks (e.g. (Cohen et al., 2015b)).

In this theoretical study, we investigate the plausibility of degenerate and stable neuromodulator circuits by focusing on the neural circuits in the midbrain which are the source of ascending pathways of two highly studied monoaminergic neuromodulators, serotonin (5-hydroxytrptamine; 5-HT) and dopamine (DA). These neuromodulators have major roles in modulating cognition, emotion and behavior, and are linked to the pathogenesis and pharmacological treatment of many common neuropsychiatric and neurological disorders (Muller and Cunningham, 2020). The majority of 5-HT-producing neurons are found in the dorsal and median raphe nuclei (DRN and MRN), while most DA-producing neurons reside in the ventral tegmental area (VTA) and substantia nigra compacta (SNc) (Muller and Cunningham, 2020).

At the functional level, DA and 5-HT are known to play important roles in reward- and punishment-based learning and decision-making (Hu, 2016). For example, there is strong evidence that DA neuronal activity signals reward prediction error (difference between predicted and actual reward outcome) to guide reinforcement learning (Doya, 2002). Specifically, DA neurons are phasically excited upon unexpected reward outcome or reward-predictive cues and inhibited upon unexpected reward omission or punishment (Schultz et al., 1997; Cohen and Uchida, 2012; Watabe-Uchida et al., 2017), although there is heterogeneity amongst DA neurons in this regard (Cohen and Uchida, 2012; Cohen et al., 2015a).

In comparison to DA neurons, DRN 5-HT neurons exhibit greater complexity in function, with studies reporting that 5-HT neuronal activity encodes both reward and punishment. For instance, it has been shown that certain 5-HT neurons (labelled “Type I”) were phasically activated only by reward predicting cues, but not punishment in a classical conditioning paradigm (Cohen et al., 2015a). Yet, in the same study, a different population of 5-HT neurons (“Type II”) signaled both expected reward and punishment with sustained elevated activity towards reward outcome (Cohen et al., 2015b). The latter study also found that, unlike DA neurons, baseline firing of Type-I 5-HT neurons was generally higher in rewarding than punishment trials, and this effect lasted across many trials, suggesting information processing over a long timescale, possibly used to perform meta-learning via learning rate modulation (Grossman et al., 2022). Similar 5-HT neuronal responses to reward and punishment were reported in other rodent (Liu et al., 2014; Li et al., 2016; Matias et al., 2017; Zhong et al., 2017) and non-human primate (Hayashi et al., 2015) studies. This differential responses of DA and 5-HT neurons to reward and punishment cannot be trivially reconciled with a simple model e.g. using two opposing neuromodulatory systems as previously proposed (Boureau and Dayan, 2010).

Other studies reveal further complexity in reward/punishment processing, specifically in the form of altered activity of non-5-HT/DA midbrain neurons. For example, DRN neurons utilizing gamma-aminobutyric (GABA) were tonically inhibited during reward-waiting with further inhibition during reward acquisition, but phasically activated by aversive stimuli (Li et al., 2016). In contrast, GABAergic neuronal activity in the VTA exhibited sustained activity upon rewarding cue onset but no response to the presence or absence of actual reward outcome (Cohen and Uchida, 2012). Further, other studies found that VTA GABAergic neuronal activity was potently and phasically activated by punishment outcome, which in turn inhibited VTA DA neuronal activity (Tan et al., 2012; Eshel et al., 2015). Another study showed that glutamatergic (Glu) neurons in the DRN reinforced instrumental behavior through VTA DA neurons (McDevitt et al., 2014).

This complexity of signaling within the DRN-VTA system in response to reward and punishment may reflect the DRN and VTA having shared afferent inputs (Watabe-Uchida et al., 2012, 2017; Ogawa et al., 2014; Pollak Dorocic et al., 2014; Beier et al., 2015; Tian et al., 2016; Ogawa and Watabe-Uchida, 2018). Another possibility is that the DRN and VTA interact with each other. Indeed, a growing number of studies have suggested that there are direct and indirect interactions among distinctive neuronal types between and within the DRN and VTA (di Giovanni et al., 2008; Tan et al., 2012; Watabe-Uchida et al., 2012; McDevitt et al., 2014; Ogawa et al., 2014; Beier et al., 2015; de Deurwaerdère and di Giovanni, 2017; Valencia-Torres et al., 2017; Xu et al., 2017; Li et al., 2019; Wang et al., 2019).

Taken together, information of reward and punishment signaled by neuronal activities within the DRN and VTA seems to be diverse, heterogeneous, distributed and mixed. Some of these signaling responses are illustrated in Fig. 1B and summarised in Supplementary Table 1. This led to the following questions (Fig. 1A). First, can these experimental findings from separate studies be reconciled and understood in terms of a single, parsimonious neural circuit model encompassing both the DRN and VTA? Second, can there be several degenerate DRN-VTA circuits, which are stable, that can signal reward and punishment information as observed in experiments? Third, if there are degenerate and stable DRN-VTA circuits, can some of them be distinguishable from others, for example, through perturbative means?

To address these questions, we developed a biologically plausible DRN-VTA computational neural circuit model based on our previous multiscale modelling framework for neuromodulators (Joshi et al., 2017; Wong-Lin et al., 2017), constrained by some of the observed neuronal signaling profiles under reward and punishment tasks. The modelling considers known direct and indirect pathways between DRN 5-HT and VTA DA neurons. Upon simulating the model under reward and punishment conditions, we found that many, but not all of the experimental findings, could be captured in a single, parsimonious DRN-VTA model. Further, several distinct model architectures could replicate the same neural circuit activity response profile, hence reflecting the possibility of degeneracy. Applying dynamical systems theory, we found that all these circuits were dynamically stable. To distinguish among these degenerated models, we simulated drug effects of DA D2-receptor based agonist and were able to distinguish sub-groups of these degenerate model architectures. Overall, our study provides theoretical support for degeneracy and stability of neural circuits of neuromodulators.

## 2 Materials and Methods

### 2.1 DRN-VTA network modelling

To develop the DRN-VTA neural circuit models, we made use of our dynamic mean-field (neuronal population-based) modelling framework (Joshi et al., 2017) for neuromodulator interactions. The modelling approach was constrained by data from known electrophysiological, neuropharmacological and voltammetry parameters (see (Joshi et al., 2017) and below). Each neural circuit model architecture investigated consisted of DRN 5-HT, VTA DA, VTA GABA, DRN GABA, and DRN Glu neuronal populations. Direct and indirect interactions among these five neuronal populations were then explored. The main aim of this work was to evaluate the plausibility of neuromodulator circuit degeneracy and stability rather than replicate every neuronal population in these brain regions.

Directed interaction between two neuronal populations, in which the source was a neuromodulator, was mathematically described by the neuromodulator neuronal population firing (rate) activity, followed by the release-and-uptake dynamics of the neuromodulator, which in turn induced certain population-averaged currents on the targeted neuronal population (Joshi et al., 2017). An induced current, *I*_*x*_, which could be effectively excitatory or inhibitory depending on experimental findings, can be described by

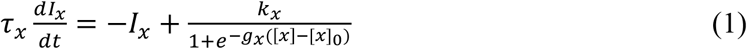

where *x* was some neuromodulator (5-HT or DA), *τ*_*x*_ the associated time constant, *k*_*x*_ some constant that determined the current amplitude, and constants *g*_*x*_ and [*x*]_0_ that controlled the slope and offset of the function on the right-hand-side of Eq. (1). The release-and-uptake dynamics for a neuromodulator *x* was described by

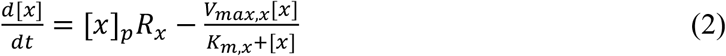

where [*x*] was the concentration of *x*, [*x*]_*p*_ the release per neural firing frequency (Joshi et al., 2017), and constants *V*_*Max,x*_ and *K*_*m,x*_ were constants determined from voltammetry measurements.

If the source of the interaction came from GABAergic (or Glu) neurons, then for simplicity, we assumed an instantaneous inhibitory (or excitatory) current-based synaptic influence on the targeted neuronal populations. This reflected the faster ionotropic receptor-based synaptic dynamics compared to metabotropic receptor-based neuromodulatory effects, while also reducing the number of free model parameters. Similarly, we also ignored the relatively faster neuronal membrane dynamics. Threshold-linear input-output function for each neuronal population was used (Joshi et al., 2017), and described by

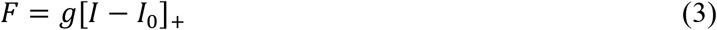

where, *F* was the neural population firing rate (output), *I* the total input current into a neural population, *g* was the slope, and *I*_0_ some threshold current, and with [*z*]_+_ = *z* if *z* ≥ 0, and 0 otherwise. Here, *I* is a function of the summed currents, including the neuromodulator-induced currents *I*_*x*_’s and ionotropic receptor-based currents (proportional to presynaptic neural firing rates). For simplicity, fast co-transmission of neurotransmitters was only considered in one modelling instance (co-release of 5-HT and glutamate via fast 5-HT3 and ionotropic receptors) based on findings by (Wang et al., 2019). From a computational modelling perspective, the DRN (Type I) 5-HT and Glu neuronal populations, which have rather similar activity profiles (Fig. 1B, 3^rd^ and 6^th^ rows) could also be effectively grouped and considered as a single 5-HT neuronal population that “co-transmit” both 5-HT and Glu to DA neurons (Wang et al., 2019).

For each model circuit architecture that we investigated, we considered separately the inclusion of either Type I or II 5-HT neurons in the circuit (Cohen et al., 2015b), and the possibility of excitatory and inhibitory projections from 5-HT to DRN Glu/GABA and DA neurons, and from DA to GABA neurons in the VTA. To allow tractability in the search for the many possible connectivity structures, the models’ connections were largely based on known experimental evidence. For example, the connections between VTA GABAergic and DRN Glu neurons were not considered as, to date, there is little experimental support. We also focused only on learned reward (with reward-predicting cue followed by reward outcome) and unexpected punishment conditions, simulated using a combination of tonic and/or phasic afferent inputs. It should be noted that we did not consider other conditions and network learning effects (e.g. (Hu, 2016)) as the main aim was to demonstrate the plausibility of DRN-VTA circuit degeneracy and stability, constrained by specific observed neuronal signalling in reward and punishment tasks. In particular, we investigated various DRN-VTA circuit architectures with network activity profiles that closely resembled those in Fig. 1B. Note also that all activity profiles in Fig. 1B were based on experimental studies except VTA Glu neural activity, which we deduced to be similar to that of DA neural activity in the reward task (McDevitt et al., 2014) and assumed to be non-responsive in the punishment task. See below for further details regarding the modelling procedures, parameters, simulations, and analyses.

### 2.2 Input-output functions of neural population firing rates

The computational models developed were based on our previous mean-field, neural population-based modelling framework for neuromodulator circuits (Joshi et al., 2017), in which the averaged concentration releases of neuromodulators (e.g. [5-HT]) were monotonic functions of the averaged firing rate of (e.g. 5-HT) neuronal populations via some neuromodulator induced slow currents. All 5 neural populations’ firing rates were described by threshold-linear functions (general form in Eq. (3)) (Jalewa et al., 2014; Joshi et al., 2017):

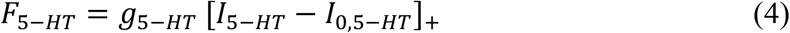

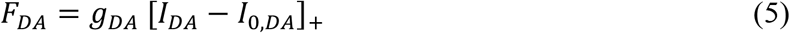

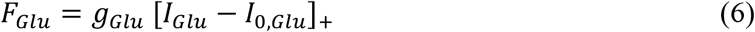

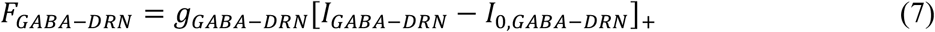

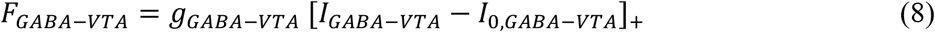

where [*z*]_+_ = *z* if *z* ≥ 0, and 0 otherwise. The threshold input values for *I*_0,*DA*_ was −10 (a.u.) for DA neurons, and *I*_0,5−*HT*_ was 0.13 (a.u.) for 5-HT neurons, to allow spontaneous activities mimicking *in vivo* conditions. 5-HT neurons had a threshold-linear function with gain value *g*_5−*HT*_ of about 1.7 times higher than that for DA neurons, and so we set their for DA and 5-HT neurons to be 0.019 and 0.033 (Hz), respectively (e.g. (Shepard and Bunney, 1991; Richards et al., 1997; Crawford et al., 2010; Wong-Lin et al., 2012; Challis et al., 2013)). For simplicity, we assumed the same current-frequency or input-output function in either tonic or phasic activity mode (Jalewa et al., 2014; Joshi et al., 2017).

### 2.3 Afferent currents and connectivity

The afferent current, *I*, for a neural population consisted of summed contributions from external excitatory inputs *I*_*ext*_ including those induced by reward or aversive stimuli, and recurrent interactions with other neural populations (see below) e.g. *I*_5−*HT,DA*_ for effective DA-induced currents in 5-HT neurons. Additionally, for a neuromodulator population, auto receptor-induced current, *I*_*auto*_, was included.

Due to limited experimental evidence, and to reduce the parameter search space, we did not consider the following connections from: (i) DRN GABA to VTA DA neurons; (ii) DRN GABA to VTA GABA neurons; (iii) VTA GABA to DRN Glu neurons; (iv, v) DRN Glu to DRN GABA neurons, and vice versa; and (vi) VTA DA to DRN Glu neurons. Then the total (population-averaged) afferent input currents to DA and 5-HT neurons were, respectively, described by

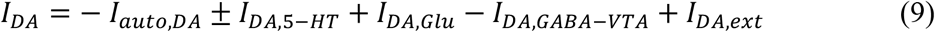

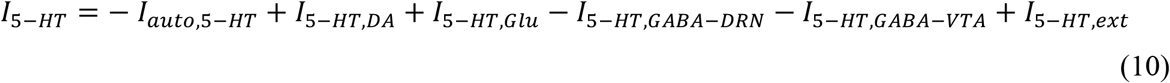

where the first terms on the right-hand sides of Eqs. (6) and (7) were autoreceptor-induced self-inhibitory currents, the second terms were the 5-HT-to-DA (labelled with subscript *DA*, 5 − *HT*) and DA-to-5-HT (with subscript 5 − *HT, DA*) interactions, the third terms were excitatory interactions from DRN Glu neurons, the fourth/fifth terms were inhibitory interactions from local DRN/VTA GABAergic neurons, and the last terms were additional external constant biased inputs from the rest of the brain and the influence of behaviorally relevant stimuli (due to rewards or punishments; see below). 5-HT neurons have a possible additional negative interaction from VTA GABA neurons (second last term on right-hand-side) (Li et al., 2019). Negative or positive sign in front of each term indicated whether the interaction was effectively inhibitory or excitatory. The ± sign indicated effectively excitatory (+) or inhibitory (−) interactions which we investigated (Figs. 3, 4 and 5), given their mixed findings in the literature (see also Eqs. (11-13)). This form of summed currents was consistent with some experimental evidence that showed different afferents modulating the tonic and phasic activation (e.g. (Floresco et al., 2003; Tian et al., 2016)).

**Figure 2.**
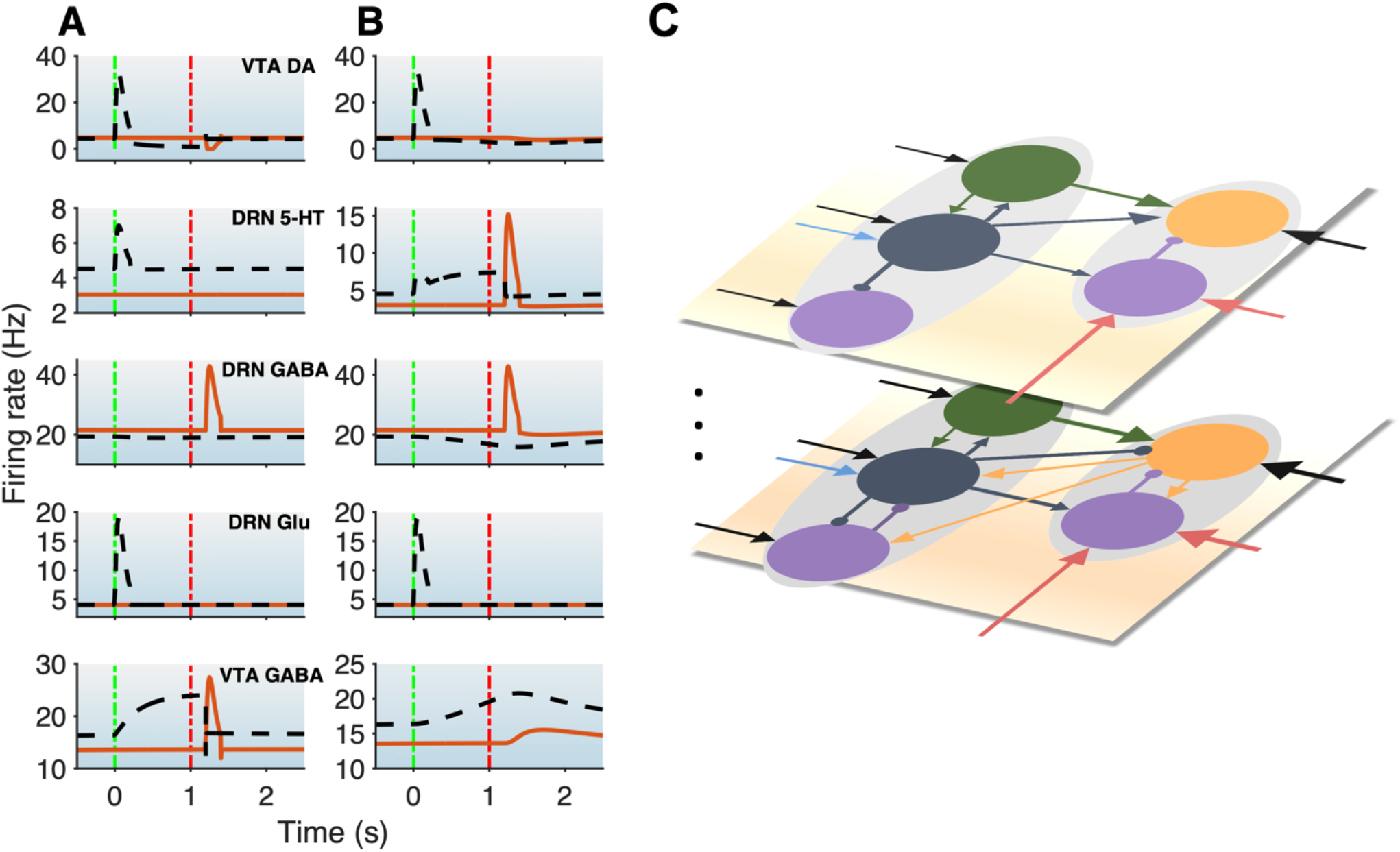
DRN-VTA model replicates signalling patterns and suggests multiple parallel circuits. **A-B)** Model with reward (black dashed lines) and punishment (orange bold lines) tasks with 5-HT neurons that are of Type I **(A)** or Type II **(B)**. This will be used as a standard activity profile template to test for degeneracy. Time label from cue onset. Green (red) vertical dashed-dotted lines: cue (outcome) onset time (as in Fig. 1B). Top-to-bottom: VTA DA, DRN 5-HT, DRN GABAergic, DRN Glu, and VTA GABAergic neural populations. **C)** Hypothesis for multiple different DRN-VTA circuits operating in parallel, which may consist of different clusters of neuronal sub-populations and different set of afferent inputs, leading to different outputs. Vertical dots denote the potential of having more than two distinctive circuits.

**Figure 3.**
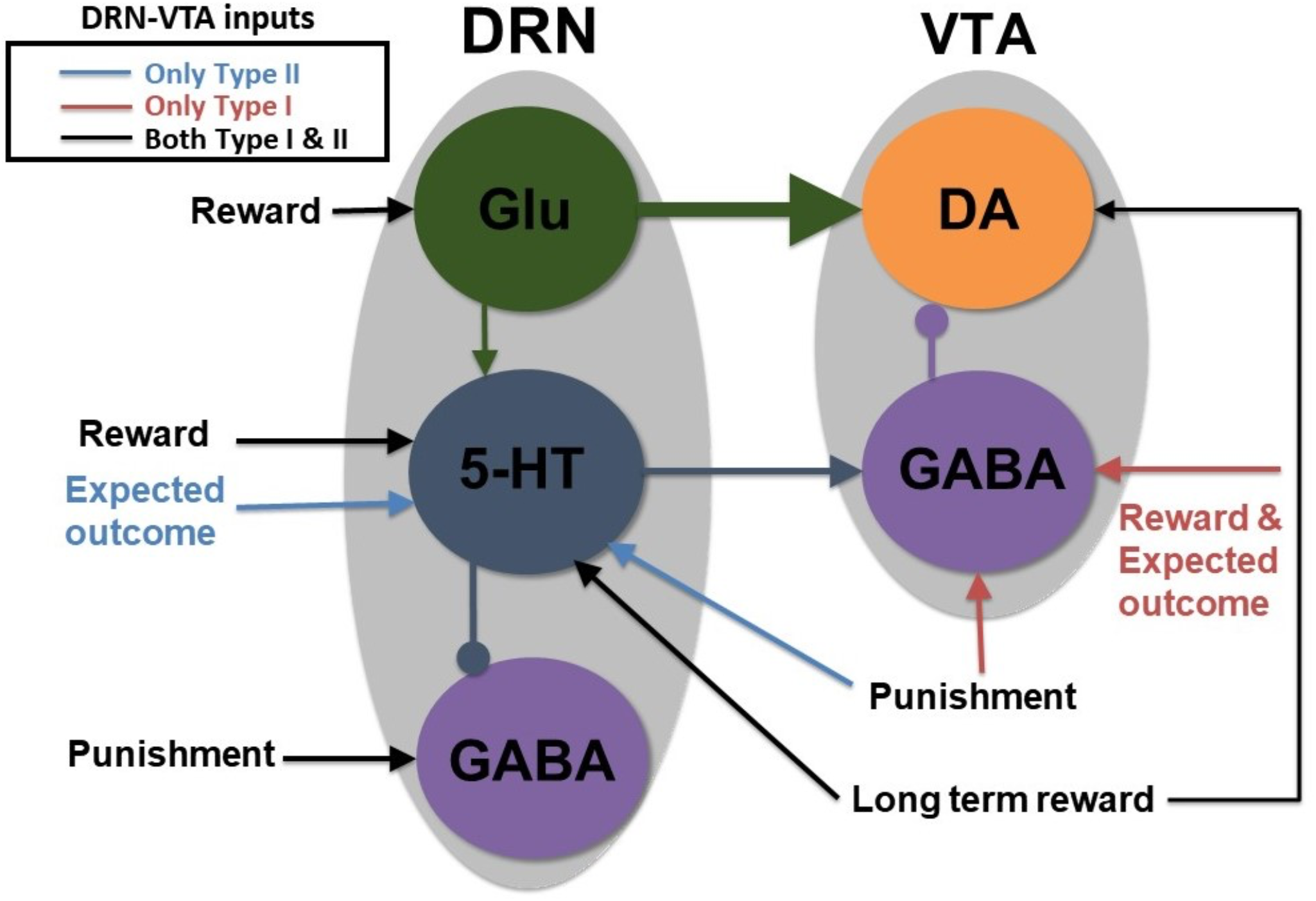
A parsimonious, sparsely connected DRN-VTA circuit model. Grey: brain region. Coloured circle: neuronal population. Legend: network’s afferent inputs. Model architecture implicitly encompasses either Type I or II 5-HT neurons with two different inputs for reward/punishment task (bright red arrows if Type I; blue arrows if Type II; black arrows denote common inputs for reward/punishment task for both Types). Circuit connections: triangular-end arrows (excitatory); circle-end arrows (inhibitory). Thicker arrows: stronger connection weights. Constant long-term reward inputs simultaneously to 5-HT and DA neurons to alter baseline activities. Sustained activity for expectation of reward outcome implemented with tonic input between cue and reward outcome. All other inputs are brief, at cue or reward/punishment outcome, producing phasic excitations/inhibitions. Note: Self-inhibitory (self-excitatory) connections within GABAergic (Glu) neurons, and auto-receptor inhibitions within 5-HT or DA neurons were implemented but not shown here (see Materials and methods, and Supplementary Information, Fig. 1). This is the most basic, sparsely connected model architecture considered.

**Figure 4.**
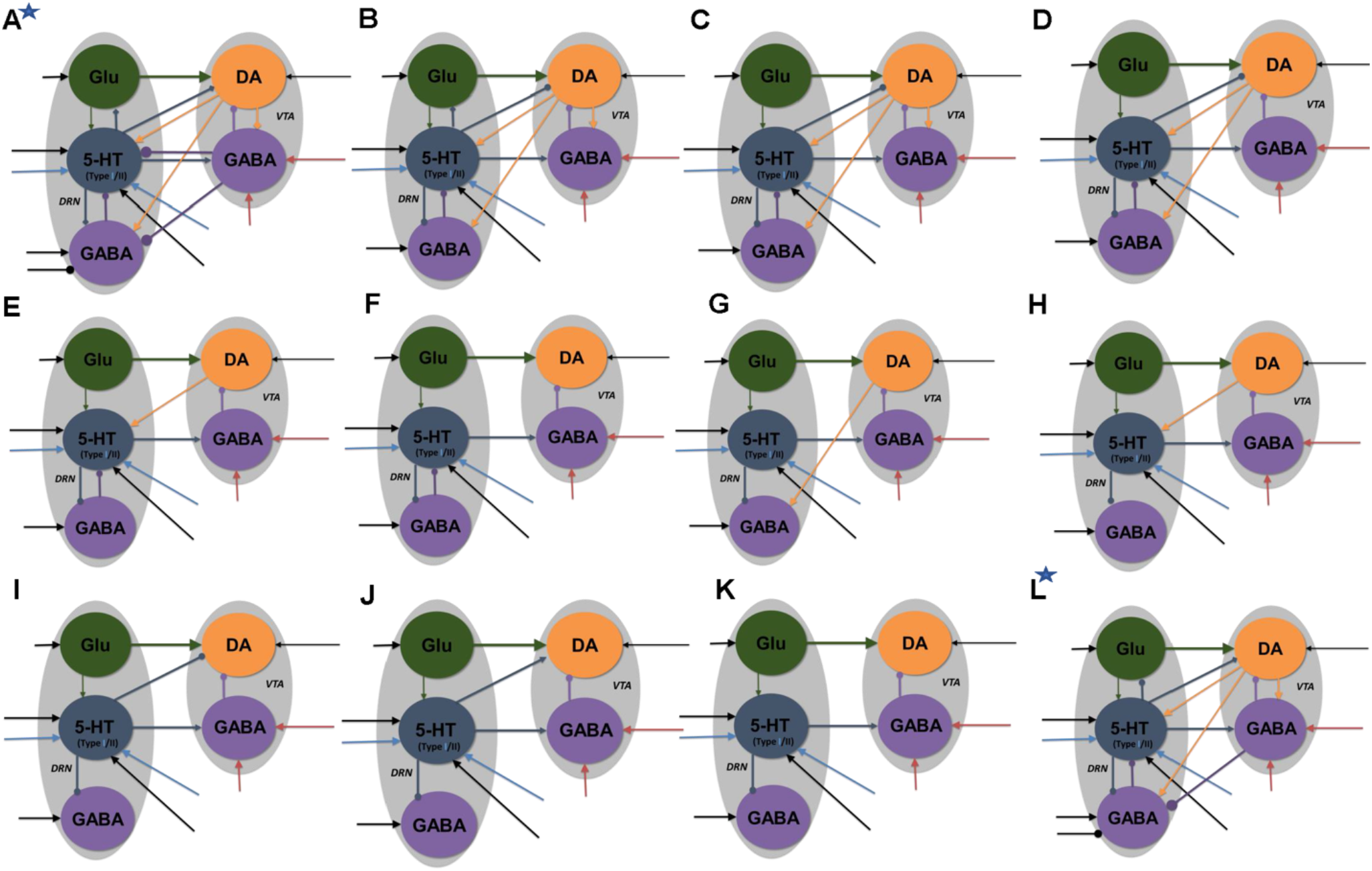
Neural circuit model architectures with similar network activity profiles. The activity profiles were similar to that in Figs. 3A and B, satisfying the inclusion criterion. Model architectures ‘A’-’K’ denote architectures of decreasing connectivity, with Fig. 3 as architecture ‘K’. Model architecture ‘L’ has an asterisk to denote that it was the only model with fast 5-HT to VTA DA connection, simulating fast 5-HT3 or Glu receptor mediated connection or their combination (co-transmission). Architectures ‘A’ and ‘L’ have additional inhibitory input to DRN GABA neurons in reward task. All labels, connections and nomenclature have the same meaning as that in Fig. 2, except that, for simplicity, the relative connection weights (thickness) are not illustrated, and the diamond-end arrows denote connections which are either excitatory or inhibitory, with both explored. Self-connectivity not illustrated (see the general example in Supplementary Information, Fig. 1 for a detailed version of model architecture ‘A’). Note: Each architecture consists of several distinctive model types (with a total of 84 types) with different 5-HT neuronal or excitatory/inhibitory connectivity types (see Supplementary Information, Tables 2 and 4, and Fig. 5).

**Figure 5.**
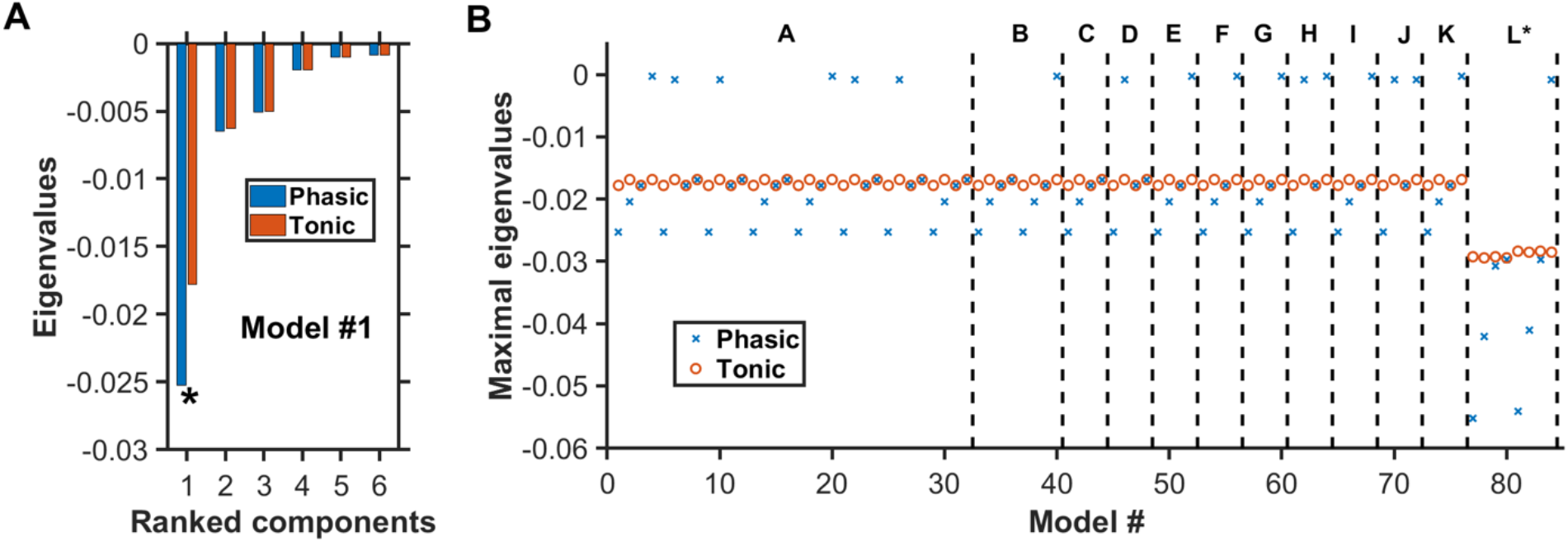
Negative real eigenvalues at steady states of degenerate models. **A)** Complete set of the real part of the eigenvalues for model #1 (Supplementary Information, Tables 2 and 4) with architecture ‘A’ in Fig. 4. Horizontal axis: Eigenvalues ranked from the largest to the smallest (magnitude wise). Blue (red): More negative eigenvalues with phasic (blue) than tonic (red) input. Asterisk: Maximal eigenvalue (largest magnitude) for each condition. **B)** For each of the 84 models, only the real part of the eigenvalue with the largest magnitude is plotted under phasic (blue cross) and tonic (red circle) input condition. Model architectures ‘A’ to ‘L’ refer to the different architectures as in Fig. 4, in which each has their own distinctive model types (e.g. different 5-HT neuronal or excitatory/inhibitory connectivity types). Eigenvalues for all model types have negative real parts, indicating dynamically stable. For most models, the eigenvalues are generally more negative during phasic than tonic activities.

Similarly, the total (population-averaged) afferent current to the glutamatergic (Glu), and VTA and DRN GABAergic neurons, can respectively be described by

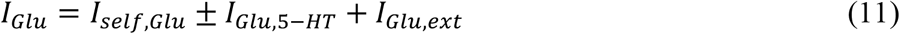

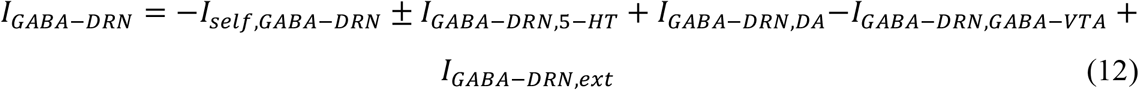

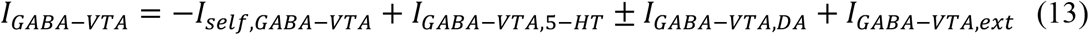

where, the subscript *Glu* denoted DRN Glu neural population, and subscripts *GABA* − *VTA* and *GABA* − *DRN* denoted GABAergic neural populations in the VTA and DRN, respectively. The subscript *self* denoted self-connection.

The averaged synaptic currents of non-5-HT/DA ionotropic glutamatergic and GABAergic neurons, namely, *I*_*DA,glu*_, *I*_5−*HT,Glu*_, *I*_*DA,GABA*−*VTA*_, *I*_5−*HT,GABA*−*VTA*_, *I*_5−*HT,GABA*−*DRN*_, *I*_*self,Glu*_, *I*_*self,GABA*−*DRN*_, *I*_*self,GABA*−*VTA*_ and *I*_*GABA*−*DRN,GABA*−*VTA*_ were typically faster than currents induced by (metabotropic) 5-HT or DA currents. Thus, we assumed the former currents to reach quasi-steady states and described (Jalewa et al., 2014) and represented by *I*_*e*/*i*_ = ± *J*_*e*/*i*_ *F*_*e*/*i*_, where the subscript *e*/*i* denoted excitatory/inhibitory synaptic current, *J* the connectivity coupling strength, *F* the presynaptic firing rate for neural population *e*/*i*, and the sign ± for excitatory or inhibitory currents. Further, dimensionless coefficients or relative connectivity weights, *W*′*s* (with values ≥ 0), were later multiplied to the above neuromodulator induced current terms (right-hand-side of terms in Eqs. (9-13); see Eqs. (20-24)). Both the *J*′*s* and *W*′*s* were allowed to vary to fit the network activity profiles of Fig. 1B within certain tolerance ranges (see below) while exploring different neural circuit architectures (Fig. 4) (see Supplementary Information, Table 2 for specific values). The self-connection weights *J*′*s* within the DRN Glu, DRN GABA and VTA GABA neurons were set at 0.5, 0.5, and 10 respectively, for all network activity’s response profiles.

Autoreceptor-induced currents were known to trigger relatively slow G protein-coupled inwardly-rectifying potassium (GIRK) currents (Tuckwell and Penington, 2014). For 5-HT1A auto-receptors, the inhibitory current *I*_5−*HT,auto*_ was described by (Ritter et al., 2008; Tuckwell and Penington, 2014; Joshi et al., 2017)

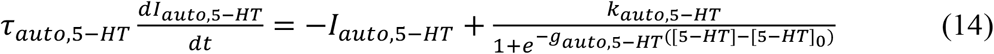

and similarly, for DA auto-receptor induced inhibitory current *I*_*auto,DA*_:

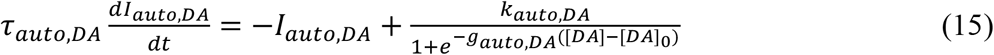

where *τ*_*auto*,5−*HT*_ was set at 500 ms (Joshi et al., 2017) and *τ*_*auto,DA*_ at 150 ms (Benoit-Marand et al., 2001; Courtney et al., 2012; Cullen and Wong-Lin, 2015). The threshold values [5 − *HT*]_0_ and [*DA*]_0_ were set at 0.1 μM. These parameters can be varied to mimic the effects of auto-receptor antagonists/agonist (Joshi et al., 2017). The gains *g*_*auto*,5−*HT*_ and *g*_*auto,DA*_ were set at 10 μM^-1^ each, and *k*_*auto*,5−*HT*_ = *k*_*auto,DA*_ = 80 a.u.. These values were selected to allow reasonable spontaneous neural firing activities and baseline neuromodulator concentration levels (see below), and similar to those observed in experiments.

Similarly, we assumed sigmoid-like influence of [5 − *HT*] ([*DA*]) on DA (5-HT) neural firing activities between the DRN and VTA populations such that the induced current dynamics could be described by (Wang and Wong-Lin, 2013; Joshi et al., 2017) :

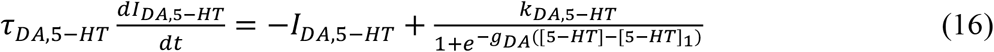

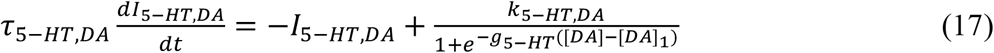

with the time constants *τ*_5−*HT,DA*_ = 1s, and *τ*_*DA*,5−*HT*_ = 1.2s (Haj-Dahmane, 2001; Aman et al., 2007). We set *k*_5−*HT,DA*_ = *k*_*DA*,5−*HT*_ = 0.03 a.u. and *g*_5−*HT*_ = *g*_*DA*_ = 20 μM^-1^, [5 − *HT*]_1_ = 0.3nM, [*DA*]_1_ = 0.1 nM such that the neural firing activities and baseline neuromodulator concentration levels were at reasonable values (see below) and similar to those in experiments (e.g. (Bunin et al., 1998; Hashemi et al., 2011)). For simplicity, we assumed Eqs. (16) and (17) to be applied equally to all targeted neural populations, but with their currents multiplied by their appropriate weights *w*′*s* (see above).

### 2.4 Release-and-reuptake dynamics of neuromodulators

The release-and-reuptake dynamics of 5-HT followed the form of a Michaelis-Menten equation (Bunin et al., 1998; Hashemi et al., 2011; Joshi et al., 2011, 2017):

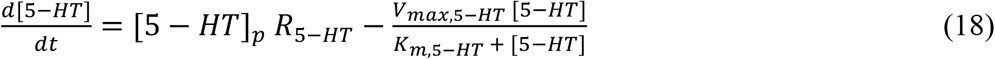

where [5 − *HT*]_*p*_ = 0.08 nM was defined as the release per firing frequency (Joshi et al., 2011, 2017; Flower and Wong-lin, 2014) (value selected to fit to reasonable baseline activities (Hashemi et al., 2011); see below), and the Michaelis-Menten constants *V*_*Max*,5−*HT*_ = 1.3 μM/s (maximum uptake rate) and *K*_*m*,5−*HT*_ = 0.17 μM (substrate concentration where uptake proceeds at half of maximum rate) were adopted from voltammetry measurements (Hashemi et al., 2011).

Similarly, the release-and-reuptake dynamics for DA was described by

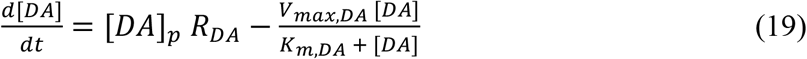

where *V*_*Max,DA*_ = −0.004 μM/s and *K*_*m,DA*_ = 0.15 μM (May et al., 1988). We set [*DA*]_*p*_ = 0.1 nM to constrain the ratio [*DA*]_*p*_/[5 − *HT*]_*p*_ = 1.25 (May et al., 1988; Bunin et al., 1998; Hashemi et al., 2011). For simplicity, we assumed Eqs. (18) and (19) to be applied equally to all targeted neural populations.

### 2.5 Reward and punishment conditions with Type I and Type II 5-HT neurons

To limit the size of the parameter search space, we focused on only the classical, fully learned reward conditioning task, and unexpected punishment task. For each simulated trial or realization within a set of conditions (reward/punishment, excitatory/inhibitory connection), we set the cue onset time at 4.5s to allow the network time to stabilize and reach steady state. The within-trial protocol for the external input current, *I*_*ext*_, was implemented as a function of time *t* as followed, depending on the simulated conditions. Note that, for simplicity, all external input currents were assumed to be excitatory, regardless of reward or punishment task, unless stated.

For reward task with Type-I 5-HT neurons: (i) constant input currents *I*_5−*HT,ext*_ and *I*_*DA,ext*_ to 5-HT and VTA DA neurons, respectively, of amplitude 50 *a. u*. to simulate long-term reward; (ii) brief 0.2s pulse *I*_*Glu,ext*_ to DRN Glu neurons of amplitude 1000 *a. u*. at 4.5s; (iii) constant step input current *I*_*GABA*−*VTA,ext*_ to VTA GABAergic neurons with amplitude 200 *a. u*. from 4.5 to 5.7s to simulate sustained activity; and (iv) no external input *I*_*GABA*−*DRN,ext*_ to DRN GABAergic neurons. For punishment task with Type-I 5-HT neurons: (i) no external input to VTA DA, 5-HT neurons and DRN Glu neurons; (ii) brief 0.2s pulse to VTA (*I*_*app*_) and DRN GABAergic (*I*_*app*5*i*_) neurons with amplitude 1000 *a. u*. at 5.7s.

For reward task with Type II 5-HT neurons: (i) constant input current to 5-HT and VTA DA neurons with amplitude 50 a.u. to simulate long-term reward; (ii) additional step input current to 5-HT neurons with amplitude 100 a.u. from 4.5 to 5.7s to simulate sustained activity; (iii) brief 0.2 s pulse to DRN Glu neurons of 1000 a.u. at 4.5 s; (iv) no external input to VTA and DRN GABAergic neurons. For punishment task with Type II 5-HT neurons: (i) no external input to VTA DA and GABAergic neurons, and DRN Glu neurons; (ii) brief 0.2s pulse to DRN GABAergic and 5-HT neurons with amplitude of 1000 a.u. at 5.7s.

In addition, for transient inputs, we have used multiplicative exponential factors exp(−*t*/*τ*) with *τ* of 50 ms to smooth out activity time courses (e.g. afferent synaptic filtering), but they do not affect the overall results. To simulate long-term, across-trial reward/punishment signaling, we assumed a higher constant excitatory input to both 5-HT and DA neurons in reward than punishment trials. When searching for the neural circuit architecture using either Type I or II 5-HT neurons, we limited ourselves as much as possible to the same internal DRN-VTA circuit structure. This also reduced the complexity of the parameter search space.

### 2.6 Baseline neural activities and inclusion criterion

We define the neural circuit activities under baseline condition (right before cue onset) to follow that in Figs. 2A-B. Namely, the baseline firing rates for 5-HT, DRN GABA, DRN Glu, DA and VTA GABA neurons in punishment task were 3.0, 21.5, 4.1, 4.8 and 13.5 Hz, and those in reward task were 4.5, 19.4, 4.1, 4.8 and 16.3 Hz, respectively (compared with Supplementary Information, Table 3). Baseline [5 − *HT*] and [*DA*] levels were constrained to be at 10 and 1.5 nM, respectively. However, it is known that these activities can vary widely across subjects, species, and studies (Supplementary Information, Table 3).

While searching for degenerate neural circuit architecture or investigating substantial changes due to simulated D_2_ agonist, we had to define acceptable ranges of neural activities to evaluate whether the variant neural circuit still behaved similarly to that of the model in Figs. 2A-B. Specifically, an inclusion criterion was set that evaluated the time-averaged percentage change in the neural population activities of any new model with respect to that of the template activity profiles (Figs. 2A-B); the time-averaged percentage change in activities of the DA, 5-HT, DRN GABA, VTA GABA, and Glu neural populations for any qualified (degenerate) circuits had to be less than 10%, 10%, 16%, 16% and 10% from those of the template activity profiles, respectively. These mean percentage changes were calculated within a 3 s time duration, from 1s before cue onset to 1s after outcome onset, encompassing both baseline and stimulus-evoked activities. Any neural circuit architecture which did not satisfy the inclusion criterion for any neural population activity was discarded when in search of degenerate neural circuits, or was highlighted (embedded in non-black region in Fig. 6) when investigating simulated D_2_ agonist (see below). As this part of the study was to proof a concept, specific values of the inclusion criterion were theoretical and not critical - the general results, i.e. overall trends, remained even if these values were altered.

**Figure 6.**
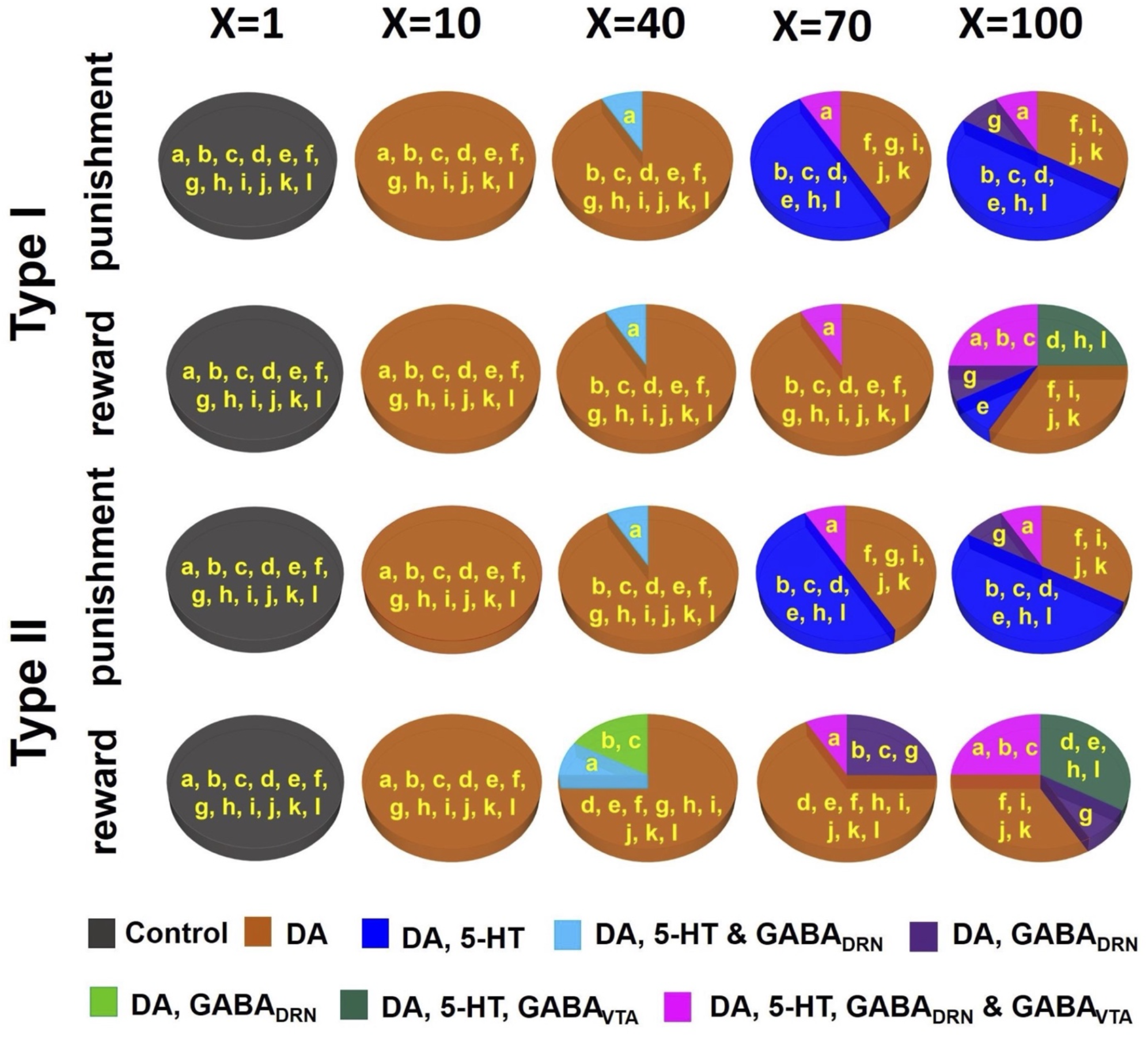
D_2_ receptor agonist can distinguish subsets of DRN-VTA neural circuits. Simulated drug administered during punishment and reward tasks with efficacy factor X increments of 1, 10, 40, 70 and 100 times. Baseline condition: X=1. Colours other than black in the pie chart denote that at least one neural population activity in specific model architecture(s) (labelled in yellow, and as in Fig. 4) has deviated (increased or decreased) beyond the inclusion criterion. Specific colours denote changes in specific neural population activities (See Supplementary Information, Tables 5-9 for details.)

### 2.7 Simulating the effects of D2 agonist

DA and 5-HT induced currents can lead to overall excitatory or inhibitory effects, depending on receptor subtype(s) and the targeted neurons. In particular, DA enhances VTA GABAergic neuronal activity via D_2_ receptors and depolarizes the membrane of 5-HT neurons (e.g. (Haj-Dahmane, 2001; Aman et al., 2007; Ludlow et al., 2009; Courtney et al., 2012; Ford, 2014)). DA also regulates the activity of other DA neurons via D_2_ auto-inhibitory receptors (Adell and Artigas, 2004). To study how D_2_ receptor mediated drugs can affect DRN-VTA architecture differently, we simulated different D_2_ agonist dosage levels simultaneously by multiplying the connection weights of D_2_ receptor mediated currents (see above; Fig. 4; orange connections in Supplementary Information, Fig. 1) by a factor ‘X’ (Fig. 6) of 10, 40, 70 and 100 times (default of X=1). Then for each dosage, we separately observed the deviation in activity profiles for each neural population with respect to the allowed range. Again, we considered the time-averaged percentage changes due to the drug to be substantial if the mean percentage change in at least one neural population activity had violated the inclusion criterion (see above, and Supplementary Information, Tables 3 and 4-8).

### 2.8 Network stability analysis

The 5 neural population firing rates (Eqs. (4-8)), when combined with their associated afferent currents (Eqs. (9-13)) with explicit parameter values and relative connectivity weights, *W*’s and *J*’s (Supplementary Information, Fig. 1 and Table 2), can be rewritten, respectively, as:

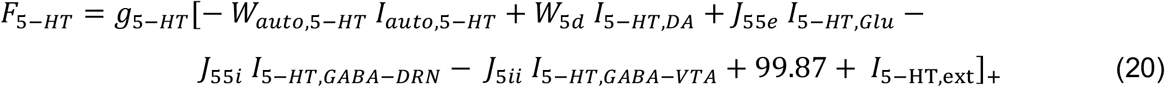

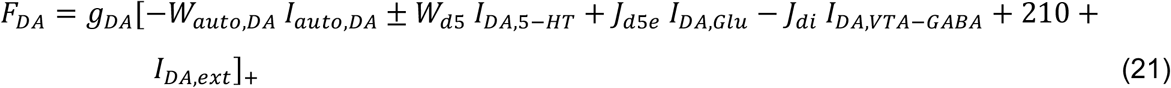

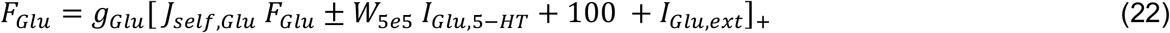

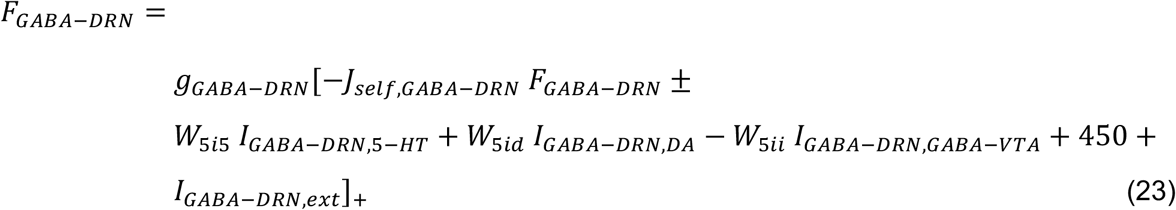

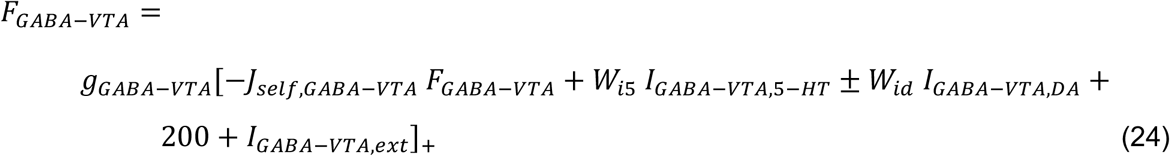

We also inserted the explicit parameter values to the dynamical equations (Eqs. (14-19)) to obtain:

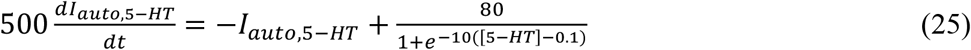

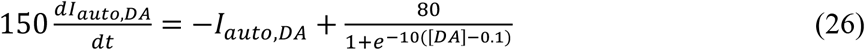

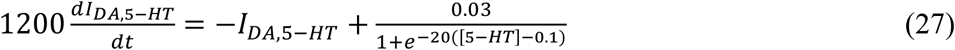

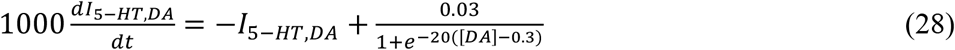

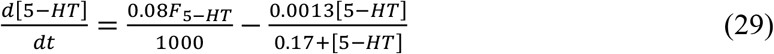

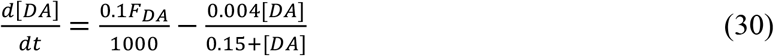

To check for network stability for each of the considered degenerate neural circuits, we first find each network’s steady state (or fixed point) by setting the rate of change for all the dynamical equations to zero, i.e 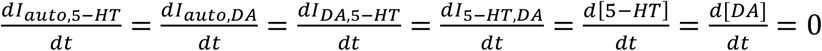, and then solving them algebraically. The solution of these equations will give the steady-state value (equilibrium point) of the system. Specifically, the currents (dynamical variables) from Eqs. (25-28) (e.g. 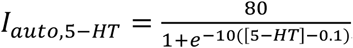) were substituted into Eqs. (20-24). See Supplementary Information, Supplementary Method, for more detailed mathematical derivation.

Using only the linear parts of the above threshold-linear functions (which were validated post-hoc), and after some algebraic manipulations, we obtained the following system of equations, in matrix form:

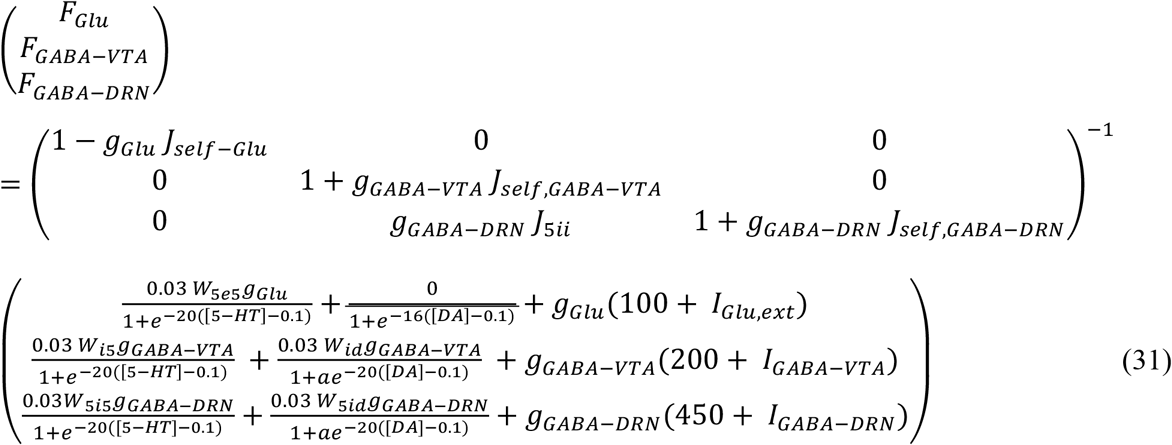

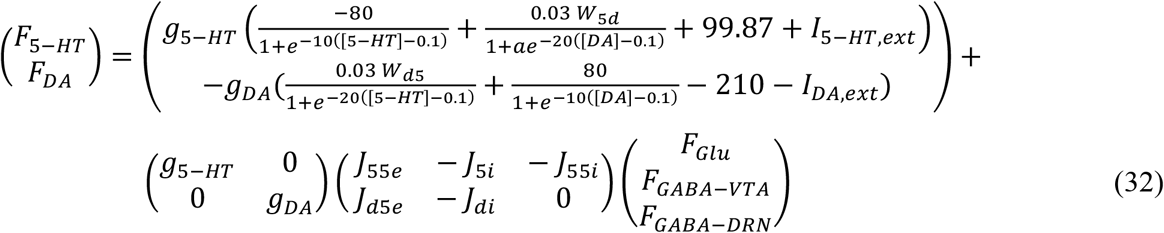

But by setting Eqs. (29) and (30) to be zero, we obtained, 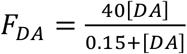 and 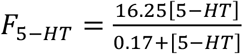 at steady state. Substituting these two into Eqns. (31) and (32), we can solve for the values of [DA] and [5-HT] at steady state.

Next, to check whether the system is stable at a steady state, we compute the 5-by-5 Jacobian matrix *M*_*Jacobian*_ (Strogatz, 2018) for the 5 dynamical equations (Eqs. (25-30)), which can be computed using partial derivatives on the right-hand-side of these equations:

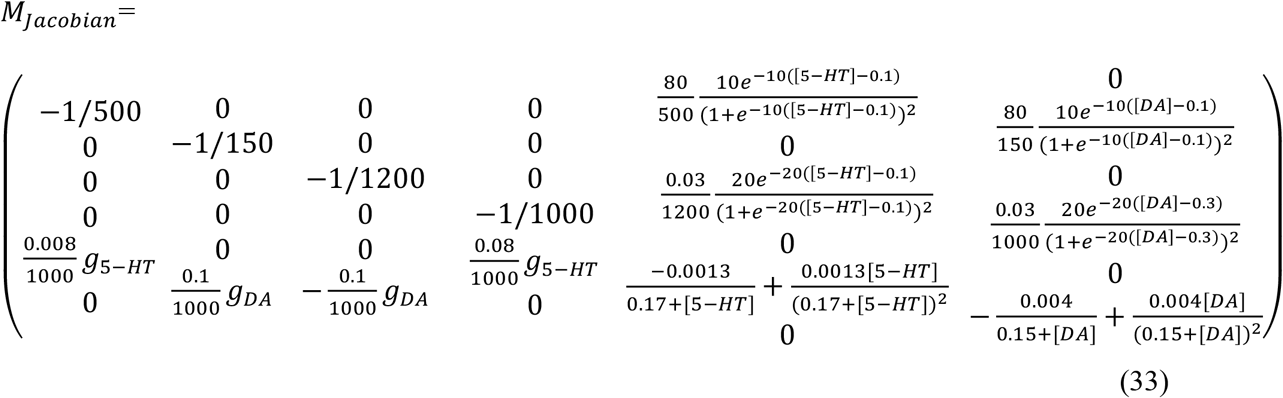

The eigenvalues of this Jacobian matrix were computed for each steady state for each model under each simulated condition (e.g. reward/punishment task). If the real parts of all the eigenvalues of the Jacobian matrix were negative at a given steady state, then the model was considered to be dynamically stable at that steady state (Strogatz, 2018).

## 3 Results

### 3.1 A DRN-VTA model can reconcile many signaling patterns

We began with a parsimonious, sparsely connected DRN-VTA model architecture and adjusted the strength of the afferent inputs and the internal connection weights of the network (Materials and Methods) such that the network activity profiles attained (Fig. 2) were readily comparable to the stereotypical profiles as illustrated in Fig. 1B. We started the simulations using a minimal network configuration, the most sparsely connected DRN-VTA circuit model investigated in this work (Fig. 3). All other subsequent architectures considered would henceforth be derived from this basic architecture. This minimal network architecture readily recapitulated many of the neuronal signalling changes in the DRN and VTA in separate experimental studies, both for reward (Fig. 2, black dashed) and punishment (Fig. 2, orange bold) tasks. The neural activity generated from the model (Figs. 2A and B) will be used as a standard activity profile template to later test for degeneracy. Deviations from this template will be later used to distinguish different subgroups of the DRN-VTA degenerate networks (see below).

In particular, in punishment task with Type I 5-HT neurons, brief excitatory inputs to GABAergic neurons in the DRN and VTA led to their punishment-based phasic activation (Fig. 2A, 3^rd^ and 5^th^ rows, orange) (Tan et al., 2012; Li et al., 2016). To replicate the activity profiles for a circuit with Type II 5-HT neurons, brief excitatory inputs to the DRN GABAergic and 5-HT neurons were implemented instead, leading to punishment-based phasic activation for both (Cohen and Uchida, 2012; Tan et al., 2012; Li et al., 2016) (Fig. 2B, 2^nd^ and 3^rd^ rows, orange). With both Type I and II 5-HT neurons, there was also phasic inhibition of VTA DA activity via VTA GABAergic neurons (Fig. 2, 1st row, orange), in line with findings from previous studies (Schultz et al., 1997; Cohen and Uchida, 2012; Tan et al., 2012; Watabe-Uchida et al., 2017). It should be noted that with Type II 5-HT neurons, the phasic activation of VTA GABAergic neurons was not as potent due to the filtering by the slow excitatory connection from 5-HT to VTA GABA neurons. Alternatively, a phasic input might instead be sent to VTA GABAergic neurons, leading to a more potent activation of the latter.

In reward task, to replicate a sustained Type II 5-HT neuronal reward signaling between cue onset and reward outcome (Fig. 2B, 2^nd^ row, black dashed) (Liu et al., 2014; Cohen et al., 2015b; Li et al., 2016), tonic excitatory input to DRN Type II 5-HT was implemented. The resulting sustained 5-HT activity led to gradual suppression of DRN GABAergic activity (Fig. 2B, 3^rd^ row, black dashed) (Liu et al., 2014; Li et al., 2016) but also gradual rise in VTA GABAergic activity regardless of reward outcome (Fig. 2B, last row, black dashed) (Cohen and Uchida, 2012; Eshel et al., 2015). The effects on these two GABAergic populations were respectively due to 5-HT’s inhibitory connection to DRN GABAergic neurons and excitatory connection to VTA GABAergic neurons. With regards to the latter, evidence of such excitatory influence via 5-HT2C receptors was reported (Valencia-Torres et al., 2017).

Excitatory connection from DRN Glu to VTA DA neurons was particularly strong in the models. This, together with no (or weak) 5-HT-to-DA connection, was consistent with previous work (McDevitt et al., 2014). In reward task, brief excitatory input to the DRN Glu neuronal population led to its reward-sensitive phasic activation (Figs. 2A-B, 4^th^ row, black dashed). Alternatively, we could have directly implemented phasic excitatory input to the VTA DA neurons. In any case, the phasic activation of DA activity was consistent with the reported DA neuronal response to a fully learned reward-predicting cue (Schultz et al., 1997; Cohen and Uchida, 2012; Watabe-Uchida et al., 2017).

Variations in some of the model parameters could still recapitulate these profiles, exhibiting model robustness (Supplementary Information, Table 2). Moreover, only modifications to the afferent inputs to DRN 5-HT and VTA GABA neurons (Materials and Methods) were needed to replicate the signalling of Type I or II 5-HT neurons (Figs. 2A and B, respectively), while maintaining the same internal connectivity structure. For example, a lack of sustained reward-based activity of Type I 5-HT activity (Fig. 2B, 2^nd^ row, black dashed) required additional external input to sustain VTA GABAergic neural activity (Fig. 2A, bottom row, black dashed; Fig. 3). It should be noted that this is assumed that the activity profiles for these non-5-HT neurons were qualitatively similar, regardless of the 5-HT neuronal types (Fig. 3).

To understand across-trial reward versus punishment effects in the model, we implemented a higher constant excitatory input into both DRN 5-HT and VTA DA neurons under reward compared to punishment conditions (Fig. 3, long black arrows). This particular model required differential inputs to 5-HT and DA neurons (Materials and methods) such that the overall tonic 5-HT neural activity was higher for reward than punishment trials, while DA neural activity remained unchanged (Fig. 2, black dashed vs orange bold lines in top two rows), again consistent with experimental observation (e.g. (Cohen et al., 2015b)). In the model, although both 5-HT and DA neurons directly received constant across-trial reward-based excitatory inputs, the indirect inhibitory pathway from 5-HT neurons through VTA GABA neurons onto VTA DA neurons nullified the overall effects on DA neurons (Figs. 2 and 3). In other words, increased firing of 5-HT neurons could be activating VTA GABAergic neurons to a level sufficient to inhibit VTA DA neurons and thereby cancelling out the net long-term reward signals (Fig. 3). Note that this is just one out of several possible models; e.g. a more complex case (with more model parameters) which we are not considering could be that the VTA DA, VTA, GABA and DRN 5-HT neurons individually receive very different inputs.

However, under these conditions, with both Type I and II 5-HT neurons, the baseline DRN and VTA GABAergic activities in the reward task were slightly different than those in the punishment task (Fig. 2, 3^rd^ and 5^th^ rows, black dashed vs orange bold), which have yet to be observed in experiments. The model’s inability to recapitulate this specific phenomenon might perhaps suggest that there are more complex features in the system, such as further division of neuronal subgroups. This also holds for other model architectures (see below, Fig. 4). For example, it could be possible that a sub-population of DRN GABAergic neurons be directly connected to 5-HT neurons (as in Fig. 1B), but not another DRN GABAergic neuronal sub-population, such that across-trial reward signal inputs are distributed differently than that of Fig. 1B. Moreover, high chemical and functional diversity amongst DRN 5-HT neurons is now well recognised (Okaty et al., 2019). Hence, it might be possible that there could exist multiple neural circuits with different circuit architectures operating in parallel, as illustrated in Fig. 2C. Another possibility could be due to mere different inputs from e.g. different sources.

In summary, we have shown that, under reward and punishment conditions, many of the observed signalling patterns in different DRN and VTA neuronal types could readily be reconciled within a single sparsely connected DRN-VTA circuit model. However, not all the signalling patterns could be captured, suggesting that multiple different neuronal sub-populations and circuits may be operating in parallel within the DRN-VTA system. Next, we shall investigate whether various different DRN-VTA neural circuits could produce the same output, i.e. be degenerate.

### 3.2 Multiple degenerate DRN-VTA circuits

To search for degenerate DRN-VTA circuit models, we evaluated various combinations of connections within and between the DRN and VTA, and where necessary, adjusted any afferent inputs (Materials and methods). We used the network activity profile template as illustrated in Figs. 2A and B as output “target” to check whether other different circuits could replicate similar activity profiles.

Given the variability of neuronal firing rates reported in the literature (Supplementary Information, Table 3), to be considered a degenerate neural circuit, we set an inclusion criterion that allowed the time-averaged percentage changes of the neural population firing activities to be within certain acceptable ranges as compared to the activities in Figs. 2A-B. Specifically, the time-averaged percentage change in activities of the DA, 5-HT, DRN GABA, VTA GABA, and Glu neural populations for any qualified (degenerate) circuits had to be less than 10%, 10%, 16%, 16% and 10% from those of the template activity profiles, respectively. These mean percentage changes were calculated within a 3 s time duration, from 1 s before cue onset to 1 s after outcome onset, encompassing both baseline and stimulus-evoked activities. Any neural circuit architecture which did not satisfy the inclusion criterion for any neural population activity was discarded. Note that here, we aimed to proof a concept and the specific values of the inclusion criterion were theoretical and not critical - the general results, i.e. overall trends, remained even if the values were altered (not shown).

Various neural circuits were created by systematic addition and modification of connections of our parsimonious, sparsely connected DRN-VTA circuit model (now presented as model architecture ‘K’ in Fig. 4). For each model architecture, model parameters were searched, in both reward and punishment tasks using both Type I and II 5-HT neurons, until the activity outputs lie within the inclusion criterion. Both excitatory and inhibitory connections from DRN 5-HT to DRN Glu/GABA and VTA DA neurons were also explored. Once the inclusion criterion was met, the model parameter values were varied to check for robustness (Supplementary Information, Table 2). This was repeated for different model architecture until we reached a highly connectivity model structure (architecture ‘A’ in Fig. 4).

Based on this extensive search process, we obtained a total of 84 different neural circuit model architectures that recapitulated the activity profiles in Figs. 2A and B. Their high-level model architectures were illustrated in Fig. 4 (see also Fig. 5, and Supplementary Information, Tables 2 and 3). For instance, model architecture ‘A’ in Fig. 4 (see Supplementary Information, Fig. 1, for detailed architecture) actually consisted of 32 distinctive models with different 5-HT neuron types and connectivity signs (Supplementary Information, Table 4). Interestingly, this architecture’s model parameters remained the same with either excitatory or inhibitory connections (Fig. 3, diamond connections; Supplementary Information, Table 2).

We observed that all models with connection from 5-HT to DA neurons to be relatively weak or not required, consistent with previous work (McDevitt et al., 2014) (but see below, (di Giovanni et al., 2008; Wang et al., 2019)). Further, a substantial number of other connections in the DRN-VTA model were identified to be redundant or relatively weak, at least in the context of the conditions investigated (Fig. 4). In particular, the relatively weak VTA DA-to-GABA connection was consistent with studies that showed either a weak or non-existent direct effect of VTA DA on VTA GABA neurons (Morales and Margolis, 2017).

With these degenerate models, we could also investigate the effects of specific DRN-VTA connectivity in which there are mixed findings or lack of evidence in the literature. For example, the influence of 5-HT on DRN GABA neurons could be excitatory or inhibitory (Hernández-Vázquez et al., 2019). In the case when this connection was inhibitory, and not excitatory, we found it to involve several degenerate circuits (model architectures ‘B’ to ‘L’). In another example, with model architecture ‘B’, when the connection from DRN 5-HT to DRN Glu neurons was inhibitory, it led to more restricted values in its connectivity strength (0-0.1 a.u.) and hence less robust than when it was excitatory (0-1 a.u.) (Supplementary Information, Table 2). It has been reported that there are mixed findings of 5-HT on VTA DA neurons (de Deurwaerdère and di Giovanni, 2017). In our simulations, we found that an inhibitory connection from 5-HT to VTA DA neurons could lead to several degenerate models (architectures ‘A’-’D’, ‘I’ and ‘L’). Note that in model ‘L’ (labelled with an extra asterisk in Fig. 4), we had also successfully simulated its fast connectivity version, mimicking either 5-HT3 receptor mediated transmission or 5-HT-glutamate co-transmission (Wang et al., 2019).

In terms of compensatory network effects, we found that upon the removal of the 5-HT to DRN GABA connection (from architecture ‘B’ to ‘C’), be it excitatory or inhibitory, the excitatory connection from DRN Glu to 5-HT neurons had to be weakened by ∼16% to satisfy the inclusion criterion. With additional removal of the connection from VTA DA to VTA GABA neurons, we found that the strength of the inhibitory connection from 5-HT to DRN GABA neurons had to be increased by ∼150% (architecture ‘D’). Interestingly, unlike other models, only the models with architectures ‘A’ and ‘L’ had to include inhibitory connection from VTA GABA to DRN GABA neurons, in which such connection was observed in e.g. (Li et al., 2019).

### 3.3 Degenerate DRN-VTA circuit models are dynamically stable

After identifying the theoretical existence of degenerate models, we used dynamical systems theory to determine whether they were dynamically stable, i.e. whether (local) perturbation from their steady states would eventually cause a return to their initial steady states (see Materials and methods). Specifically, the stability of each neural circuit could be determined by first finding the possible steady state(s) (i.e. fixed point(s)). This was achieved by setting all the dynamical (differential) equations to zero and finding the algebraic solutions for the dynamical variables (Materials and methods). Then the eigenvalues of the system’s Jacobian matrix at the steady states were computed (see Materials and methods for mathematical derivation of the steady states and the Jacobian matrix). For a neural circuit to be dynamically stable, the real part of all the eigenvalues associated with the steady state has to be negative. This was exactly what we found for all the identified degenerate neural circuits in Fig. 4 (Fig. 5; Materials and methods), with no imaginary part in the eigenvalues.

Fig. 5A displayed the complete set of the real part of the eigenvalues for model #1 with architecture ‘A’ in Fig. 4. This model had inhibitory connections from VTA DA to VTA GABA neurons and from 5-HT to VTA DA/Glu neurons, using Type I 5-HT neurons and under punishment conditions (Supplementary Information, Table 4). It was observed that the eigenvalues with phasic input (blue) were generally larger, magnitude wise, than those with tonic input (red). This was more pronounced for the eigenvalues with the largest magnitude (maximal eigenvalues) (e.g. see Fig. 5A). Importantly, all the eigenvalues were negative, indicating a dynamically stable network model even in the presence of additional phasic stimulus input.

We repeated the analysis for all 84 models, under both phasic and tonic input conditions. This analysis was presented in Fig. 5B only for the maximal eigenvalues (red circles and blue crosses). The non-maximal eigenvalues had similar trends across the other models. In general, with phasic activities (blue crosses), the models were more stable than with tonic activities (red circles). However, during phasic activations, there were 18 models with rather small (close to zero) eigenvalues (magnitude wise), albeit still negative. Hence, close to instability. This was not observed for tonic activations, where the (most negative) eigenvalues were found to hover within a small range of values (−0.017 to -0.016), except models with architecture ‘L’ (∼-0.03). In fact, the latter models, which were the only ones with a fast 5-HT-to-DA connection (Fig. 5B, models #77-84), were the most stable under both phasic and tonic conditions. Further, there was no difference identified between the excitatory (models #77-80) and inhibitory (models #81-84) connections. Moreover, most of their eigenvalues in phasic condition were substantially more negative than their tonic counterparts.

### 3.4 D2 mediated drugs can distinguish some degenerate DRN-VTA circuits

Given the large number of degenerate and stable DRN-VTA circuits identified, how could one distinguish among at least some of them? To address this, we investigated the neural circuit responses to simulated dopaminergic D_2_ receptor agonist due to the extensive D_2_ receptor mediated connectivity within the degenerate DRN-VTA circuits (Fig. 4; Supplementary Information, Fig. 1). In particular, in the degenerate models, D_2_ receptor mediated connections involved those from VTA DA neurons to DRN 5-HT, DRN GABA and VTA GABA neurons, and also the self-inhibitory connection (D2 auto-receptor-mediated) of DA neurons (see Materials and methods for references). To mimic the effects in the model of D_2_ receptor agonist, we gradually increased the strengths equally on the connections mediated by D_2_ receptors (Supplementary Information, Fig. 1, connections emanating from DA neurons) by some factor (X) and observed how the network activity changed.

As we gradually increased the strengths of these specific sets of connection, subsets of the degenerate models gradually behaved differently from the activity profile template in Figs. 2A and B (Fig. 6A, model architectures embedded in non-black regions), i.e. not satisfying the inclusion criterion, thus, possibly allowing us to distinguish them. (See Supplementary Information, Tables 5-9 for details.) When at low dose with ten-fold (X=10) increase in D_2_ mediated connections, all models showed substantial changes to their DA activities, beyond the inclusion criterion, regardless of reward/punishment condition and 5-HT neuronal type composition. Hence, they could not be distinguished at this dosage.

For moderately higher D_2_ agonist doses (X=40 and 70), models with architecture ‘A’ can be distinguished from others, with substantial alteration to the DRN GABA and 5-HT activities (Fig. 6A, light blue). Models with architectures ‘B’ and ‘C’ were also beginning to reveal substantial activity changes in reward condition with Type II 5-HT neurons. Further, with an increased factor of 70 under punishment condition, model with architectures ‘D’, ‘E’, ‘H’ and ‘L’ could be distinguished from others with additional changes to their 5-HT activities. With a factor of 100, a maximum of 5 subsets of model architectures could be distinguished based on activity enhancement of different combination of neuronal types. An interesting observation we found was that for the same composition of 5-HT neuronal type (I or II), the subsets of distinguishable neural circuits were different between reward and punishment tasks.

Overall, our drug simulation predicted that a gradual increment of the level of D_2_ receptor activation could lead to differential enhancements of firing rate activities that could be used to distinguish subsets of the degenerate DRN-VTA circuits.

## 4 Discussion

In this work, we have shown that neural circuits which are sources of neuromodulators can be degenerate. In particular, the work focuses on computationally modelling and analysing DRN-VTA circuits, as the DRN and VTA share structural and functional relationships among their constituent neuronal types. In particular, these circuits are involved in the regulation of several cognitive, emotional and behavioural processes, and implicated in many common and disabling neuropsychiatric conditions (Muller and Jacobs, 2009). Moreover, their neuronal signalling in reward and punishment tasks are well studied (Hu, 2016).

In this work, we developed biologically based mean-field computational models of the DRN-VTA circuit (Fig. 3) with several neuronal types, and tested them under classic conditions of (learned) reward and (unexpected) punishment. The modelling was partially constrained by known connectivity within and between the DRN and VTA regions, and their inputs from multiple other brain regions, including mixed combinations of inputs (Watabe-Uchida et al., 2012, 2017; Ogawa et al., 2014; Pollak Dorocic et al., 2014; Beier et al., 2015; Tian et al., 2016; Ogawa and Watabe-Uchida, 2018). Our work demonstrated that degenerate and stable DRN-VTA neural circuits are theoretically plausible.

We found that a parsimonious, sparsely connected version of the DRN-VTA model could reconcile many of the diverse phasic and tonic neural signalling events reported in the DRN and VTA in punishment and reward tasks observed across separate experimental studies (Figs. 1 and 2A-B). This model was evaluated using Type I and Type II 5-HT neurons in the DRN as defined electrophysiologically in a previous study (Cohen et al., 2015b). In the case of Type II 5-HT neurons in reward condition, the model suggested that sustained 5-HT neuron activity between cue and reward outcome (Cohen et al., 2015b)would lead to the gradual inhibition of DRN GABA neuron activity and enhancement of VTA GABA neuron activity (Cohen and Uchida, 2012; Li et al., 2016). The sparsely connected model could also reproduce experimental observations (Cohen et al., 2015b) of an increase in baseline firing of Type I 5-HT neurons across several trials in the rewarding task, without similar effects on VTA DA neurons, or in the punishment task (Fig. 2). Thus, this specific model suggested that slow, across-trial reward-based excitatory inputs could potentially be directly targeted to both DRN 5-HT and VTA DA neurons, and that inhibitory 5-HT to GABA to DA connectivity could cancel out the effects of the direct input to DA neurons, rendering only long timescale changes to the baseline activity of 5-HT neurons but not DA neurons (Fig. 3). It should however be noted that this is just one possible theoretical speculation offered out of many. For instance, another trivial possibility would be simply to have different separate inputs to DRN 5-HT and VTA DA and GABA neurons.

Despite the success of readily recapitulating the main observed phenomena, this relatively simple model was unable to capture some specific activity profiles with Type I 5-HT neurons, namely, the differential baseline activities of VTA GABA neurons and DRN GABA neurons between reward and punishment conditions. We later found that this was the case even for the more complex neural circuit models. Thus, perhaps additional neuronal populations and DRN-VTA circuits could be operating in parallel (Fig. 2C) or it could be just be these neurons receive different inputs from e.g. different sources. Such parallel circuits could be validated experimentally in the future, for example, using gene-targeting of specific DRN and VTA neuron subtypes and projections. Indeed, there is increasing evidence that DRN 5-HT neurons are more chemically diverse than previously expected, and that there is a high level of functional diversity in output pathways of the DRN and VTA (Watabe-Uchida et al., 2012, 2017; Wong-Lin et al., 2012; Ogawa et al., 2014; Pollak Dorocic et al., 2014; Weissbourd et al., 2014; Beier et al., 2015; Fernandez et al., 2016; Tian et al., 2016; Morales and Margolis, 2017; Zhou et al., 2017; Ogawa and Watabe-Uchida, 2018; Ren et al., 2018; Okaty et al., 2019). Further features will need to be incorporated including additional pathways and mechanisms as they are being uncovered, and this could include VTA glutamatergic neurons (McGovern et al, 2021). Clearly, the number of possible connections and models will be increased.

To demonstrate degeneracy in the DRN-VTA system, we showed that several variants of the DRN-VTA circuit model could readily recapitulate the same neuronal signalling profiles, with only occasional slight changes made to their afferent inputs (Fig. 4). Our results also suggest experimental studies on the DRN and/or VTA system, e.g. using tracing methods, to search for more than one neural circuit configuration.

Consistent with previous work (McDevitt et al., 2014), the degenerate models showed relatively weaker direct connection from DRN 5-HT to VTA DA neurons as compared to the connection from DRN Glu to VTA DA neurons. However, previous studies had also demonstrated a direct influence of DRN 5-HT on VTA DA neuron activity (e.g. (de Deurwaerdère and di Giovanni, 2017)). More recent work has shown that DRN 5-HT terminals in the VTA co-release glutamate and 5-HT, eliciting fast excitation (via ionotropic receptors) onto VTA DA neurons and increased DA release in the nucleus accumbens to facilitate reward (Wang et al., 2019). Hence, we also incorporated a model with fast 5-HT-to-DA connection (model architecture ‘L*’ in Fig. 4) and found such circuit to be plausible in terms of capturing the stereotypical reward and punishment signalling.

Next, we applied dynamical systems theory and showed that all the found degenerate circuits were dynamically stable (Fig. 5). Interestingly, we found that model architecture ‘L*’ with fast 5-HT-to-DA connection was dynamically more stable than all other architectures investigated (Fig. 5B). Future computational modelling work could explore the effects of co-transmission of neurotransmitters on neural circuit degeneracy and functioning and using more biologically realistic spiking neuronal network models across multiple scales, e.g. (Wong-Lin et al., 2012, 2017).

Finally, we simulated the gradual increase in dopaminergic D_2_ receptor activation, due to extensive D_2_ mediated connections in the investigated circuits. This was done by increasing the connection strengths emanating from the VTA DA neurons. This allowed us to distinguish subsets of the degenerate DRN-VTA circuits by identifying substantial and differential deviations in specific neural populations’ activities (Fig. 6). Interestingly, we found that for the same composition of 5-HT neuronal type (I or II), the subsets of distinguishable DRN-VTA circuits were different between reward and punishment tasks (Fig. 6A). Future experimental work could verify this model prediction.

As neuromodulators can selectively regulate degenerate neural circuits (Marder et al., 2014b; Cropper et al., 2016), our theoretical work showed that even if the neuromodulator sources vary in physical and functional forms, the effects of neuromodulation on targeted brain areas may remain the same. For example, the co-modulatory effects of DA and 5-HT on prefrontal cortical rhythms (Wang and Wong, 2013) may be robust to specific interactions between the VTA DA and DRN 5-HT neurons. From a more general perspective, our computational modelling and analytical framework could be applied to the study of degeneracy and stability of neural circuits involving the interactions of other neuromodulators such as norepinephrine/noradrenaline (e.g. Joshi et al., (2017)). It should be also noted that in the development of the models, we had resorted to a minimalist approach by focusing only on sufficiently simple neural circuit architectures that could replicate experimental observations. Future modelling work may investigate the relative relevance of these connections with respect to larger circuits involving cortical and subcortical brain regions across multiple scales, especially during adaptive learning e.g. Wong-Lin et al. (2017); Zhou et al. (2018).

Taken together, our computational modelling and analytical work suggests the existence of degeneracy and stability in DRN-VTA circuits. Some of these degenerate circuits can be dynamically more stable than others, and subsets of the degenerate circuits can be distinguished through pharmacological means. Importantly, our work opens up a new avenue of investigation on the existence, robustness and stability of neuromodulatory circuits, and has important implication on the stable and robust maintenance of neuromodulatory functions.

## Data Availability Statement

Simulations were performed using MATLAB with forward Euler numerical integration method on the dynamical (ordinary differential) equations (see above). A simulation time step of 1 ms was used and smaller time steps did not affect the results. MATLAB were used for analyses of network stability and sensitivity. Source codes and generated data are available at https://github.com/ckbehera/degeneracy.

## Author Contributions

C.K.B. and K.W.-L. conceptualized and designed the study. C.K.B., A.J., D.-H.W. and K.W.-L. acquired the data and conducted analyses. K.W.-L. supervised the study. T.S. provided guidance on the study. C.K.B., A.J. and K.W.-L. wrote the first draft of the manuscript. All authors interpreted the data and revised the manuscript.

## References

Adell, A., and Artigas, F. (2004). The somatodendritic release of dopamine in the ventral tegmental area and its regulation by afferent transmitter systems. Neurosci Biobehav Rev 28, 415–431. doi: 10.1016/j.neubiorev.2004.05.001.

Aman, T. K., Shen, R.-Y., and Haj-Dahmane, S. (2007). D2-like dopamine receptors depolarize dorsal raphe serotonin neurons through the activation of nonselective cationic conductance. The Journal of Pharmacology and Experimental Therapeutics 320, 376–385. doi: 10.1124/jpet.106.111690.

Beier, K. T., Steinberg, E. E., DeLoach, K. E., Xie, S., Miyamichi, K., Schwarz, L., et al. (2015). Circuit Architecture of VTA Dopamine Neurons Revealed by Systematic Input-Output Mapping. Cell 162, 622–634. doi: 10.1016/j.cell.2015.07.015.

Benoit-Marand, M., Borrelli, E., and Gonon, F. (2001). Inhibition of dopamine release via presynaptic D_2_ receptors: time course and functional characteristics in vivo. J Neurosci 21, 9134–9141.

Boureau, Y. L., and Dayan, P. (2010). Opponency Revisited: Competition and Cooperation Between Dopamine and Serotonin. Neuropsychopharmacology 2011 36:1 36, 74–97. doi: 10.1038/npp.2010.151.

Bunin, M. A., Prioleau, C., Mailman, R. B., and Wightman, R. M. (1998). Release and Uptake Rates of 5-Hydroxytryptamine in the Dorsal Raphe and Substantia Nigra Reticulata of the Rat Brain. J Neurochem 70, 1077–1087.

Challis, C., Boulden, J., Veerakumar, A., Espallergues, J., Vassoler, F. M., Pierce, R. C., et al. (2013). Raphe GABAergic neurons mediate the acquisition of avoidance after social defeat. Journal of Neuroscience 33, 13978–13988.

Cohen, J., and Uchida, N. (2012). Neuron-type specific signals for reward and punishment in the ventral tegmental area. Nature 482, 85–88. doi: 10.1038/nature10754.Neuron-type.

Cohen, J. Y., Amoroso, M. W., and Uchida, N. (2015a). Serotonergic neurons signal reward and punishment on multiple timescales. Elife 2015, 1–25. doi: 10.7554/eLife.06346.

Cohen, J. Y., Amoroso, M. W., and Uchida, N. (2015b). Serotonergic neurons signal reward and punishment on multiple timescales. Elife 4, e06346.

Courtney, N. A., Mamaligas, A. A., and Ford, C. P. (2012). Species Differences in Somatodendritic Dopamine Transmission Determine D2-Autoreceptor-Mediated Inhibition of Ventral Tegmental Area Neuron Firing. The Journal of Neuroscience 32, 13520 LP–13528. Available at: http://www.jneurosci.org/content/32/39/13520.abstract.

Crawford, L. K., Craige, C. P., and Beck, S. G. (2010). Increased intrinsic excitability of lateral wing serotonin neurons of the dorsal raphe: a mechanism for selective activation in stress circuits. J Neurophysiol 103, 2652–2663. doi: 10.1152/jn.01132.2009.

Cropper, E. C., Dacks, A. M., and Weiss, K. R. (2016). Consequences of degeneracy in network function. Curr Opin Neurobiol 41, 62–67. doi: 10.1016/j.conb.2016.07.008.

Cullen, M., and Wong-Lin, K. (2015). Integrated dopaminergic neuronal model with reduced intracellular processes and inhibitory autoreceptors. IET Systems Biology 9, 245–258. doi: 10.1049/iet-syb.2015.0018.

de Deurwaerdère, P., and di Giovanni, G. (2017). Serotonergic modulation of the activity of mesencephalic dopaminergic systems: Therapeutic implications. Progress in Neurobiology 151, 175–236. doi: 10.1016/j.pneurobio.2016.03.004.

di Giovanni, G., di Matteo, V., Pierucci, M., and Esposito, E. (2008). Serotonin–dopamine interaction: electrophysiological evidence. Progress in Brain Research 172, 45–71. doi: 10.1016/S0079-6123(08)00903-5.

Doya, K. (2002). Metalearning and neuromodulation. Neural Networks 15, 495–506. doi: 10.1016/S0893-6080(02)00044-8.

Edelman, G. M., and Gally, J. A. (2001). Degeneracy and complexity in biological systems. Proc Natl Acad Sci U S A 98, 13763–13768. doi: 10.1073/pnas.231499798.

Eshel, N., Bukwich, M., Rao, V., Hemmelder, V., Tian, J., and Uchida, N. (2015). Arithmetic and local circuitry underlying dopamine prediction errors. Nature 525, 243–246. doi: 10.1038/nature14855.

Fernandez, S. P., Cauli, B., Cabezas, C., Muzerelle, A., Poncer, J. C., and Gaspar, P. (2016). Multiscale single-cell analysis reveals unique phenotypes of raphe 5-HT neurons projecting to the forebrain. Brain Structure and Function 221, 4007–4025. doi: 10.1007/S00429-015-1142-4/FIGURES/7.

Floresco, S. B., West, A. R., Ash, B., Moore, H., and Grace, A. A. (2003). Afferent modulation of dopamine neuron firing differentially regulates tonic and phasic dopamine transmission. Nature Neuroscience 6, 968. Available at: http://dx.doi.org/10.1038/nn1103.

Flower, G., and Wong-lin, K. (2014). Reduced Computational Models of Serotonin. 61, 1054–1061.

Ford, C. P. (2014). The role of D2-autoreceptors in regulating dopamine neuron activity and transmission. Neuroscience 282, 13–22.

Griffiths, D.J., and Schroeter, D.F. (2018). Introduction to quantum mechanics. Third edition. Cambridge University Press.

Grossman, C. D., Bari, B. A., and Cohen, J. Y. (2022). Serotonin neurons modulate learning rate through uncertainty. Current Biology 32, 586-599.e7. doi: 10.1016/J.CUB.2021.12.006.

Haj-Dahmane, S. (2001). D2-like dopamine receptor activation excites rat dorsal raphe 5-HT neurons in vitro. Eur J Neurosci 14, 125–134.

Hashemi, P., Dankoski, E. C., Wood, K. M., Ambrose, R. E., and Wightman, R. M. (2011). In vivo electrochemical evidence for simultaneous 5-HT and histamine release in the rat substantia nigra pars reticulata following medial forebrain bundle stimulation. J Neurochem 118, 749–759.

Hayashi, K., Nakao, K., and Nakamura, K. (2015). Appetitive and aversive information coding in the primate dorsal raphe nucleus. Journal of Neuroscience 35, 6195–6208.

Hernández-Vázquez, F., Garduño, J., and Hernández-López, S. (2019). GABAergic modulation of serotonergic neurons in the dorsal raphe nucleus. Reviews in the Neurosciences 30, 289–303.

Hocking, D. R., Bradshaw, J. L., and Fielding, J. (Eds.). (2019). Degenerative disorders of the brain. Routledge.

Hu, H. (2016). Reward and Aversion. http://dx.doi.org/10.1146/annurev-neuro-070815-014106 39, 297–324. doi: 10.1146/ANNUREV-NEURO-070815-014106.

Jalewa, J., Joshi, A., McGinnity, T. M., Prasad, G., Wong-Lin, K., and Hölscher, C. (2014). Neural Circuit Interactions between the Dorsal Raphe Nucleus and the Lateral Hypothalamus: An Experimental and Computational Study. PLOS ONE 9, 1–16. doi: 10.1371/journal.pone.0088003.

Joshi, A., Wong-Lin, K., McGinnity, T. M., and Prasad, G. (2011). A mathematical model to explore the interdependence between the serotonin and orexin/hypocretin systems. in 2011 Annual International Conference of the IEEE Engineering in Medicine and Biology Society, 7270–7273. doi: 10.1109/IEMBS.2011.6091837.

Joshi, A., Youssofzadeh, V., Vemana, V., McGinnity, T. M., Prasad, G., and Wong-Lin, K. (2017). An integrated modelling framework for neural circuits with multiple neuromodulators. Journal of The Royal Society Interface 14, 20160902. doi: 10.1098/rsif.2016.0902.

Levitan, I. B. (1987). Neuromodulation: The Biochemical Control of Neuronal Excitability. Oxford University Press, USA.

Li, Y., Li, C.-Y., Xi, W., Jin, S., Wu, Z.-H., Jiang, P., et al. (2019). Rostral and Caudal Ventral Tegmental Area GABAergic Inputs to Different Dorsal Raphe Neurons Participate in Opioid Dependence. Neuron 101, 748-761.e5. doi: 10.1016/j.neuron.2018.12.012.

Li, Y., Zhong, W., Wang, D., Feng, Q., Liu, Z., Zhou, J., et al. (2016). Serotonin neurons in the dorsal raphe nucleus encode reward signals. Nat Commun 7, 1–15.

Liu, Z., Zhou, J., Li, Y., Hu, F., Lu, Y., Ma, M., et al. (2014). Dorsal raphe neurons signal reward through 5-HT and glutamate. Neuron 81, 1360–1374. doi: 10.1016/j.neuron.2014.02.010.

Ludlow, K. H., Bradley, K. D., Allison, D. W., Taylor, S. R., Yorgason, J. T., Hansen, D. M., et al. (2009). Acute and chronic ethanol modulate dopamine D2-subtype receptor responses in ventral tegmental area GABA neurons. Alcoholism: Clinical and Experimental Research 33, 804–811.

Marder, E. (2012). Neuromodulation of neuronal circuits: back to the future. Neuron 76, 1–11.

Marder, E., O’Leary, T., and Shruti, S. (2014a). Neuromodulation of circuits with variable parameters: single neurons and small circuits reveal principles of state-dependent and robust neuromodulation. Annu Rev Neurosci 37, 329–346.

Marder, E., O’Leary, T., and Shruti, S. (2014b). Neuromodulation of circuits with variable parameters: single neurons and small circuits reveal principles of state-dependent and robust neuromodulation. Annu Rev Neurosci 37, 329–346. doi: 10.1146/annurev-neuro-071013-013958.

Matias, S., Lottem, E., Dugué, G. P., and Mainen, Z. F. (2017). Activity patterns of serotonin neurons underlying cognitive flexibility. Elife 6. doi: 10.7554/ELIFE.20552.

May, L. J., Kuhr, W. G., and Wightman, R. M. (1988). Differentiation of Dopamine Overflow and Uptake Processes in the Extracellular Fluid of the Rat Caudate Nucleus with Fast-Scan In Vivo Voltammetry. Journal of Neurochemistry 51, 1060–1069. doi: 10.1111/J.1471-4159.1988.TB03069.X.

McDevitt, R. A., Tiran-Cappello, A., Shen, H., Balderas, I., Britt, J. P., Marino, R. A. M., et al. (2014). Serotonergic versus non-serotonergic dorsal raphe projection neurons: differential participation in reward circuitry. Cell Reports 8, 1857–1869. doi: 10.1016/j.jacc.2007.01.076.White.

McGovern, Dillon J., Polter Abigail M., Root David H. (2021). Neurochemical Signaling of Reward and Aversion to Ventral Tegmental Area Glutamate Neurons. J. Neurosci. 41, 5471–5486, doi: 10.1523/JNEUROSCI.1419-20.2021.

Morales, M., and Margolis, E. B. (2017). Ventral tegmental area: cellular heterogeneity, connectivity and behaviour. Nature Reviews Neuroscience 18, 73.

Muller, C. P., and Cunningham, K. A. (2020). Handbook of the behavioral neurobiology of serotonin. Academic Press.

Muller, C. P., and Jacobs, B. (2009). Handbook of the Behavioral Neurobiology of Serotonin, Handbook of Behavioral Neuroscience.

Ogawa, S. K., Cohen, J. Y., Hwang, D., Uchida, N., and Watabe-Uchida, M. (2014). Organization of monosynaptic inputs to the serotonin and dopamine neuromodulatory systems. Cell Rep 8, 1105–1118.

Ogawa, S. K., and Watabe-Uchida, M. (2018). Organization of dopamine and serotonin system: Anatomical and functional mapping of monosynaptic inputs using rabies virus. Pharmacology Biochemistry and Behavior 174, 9–22.

Okaty, B. W., Commons, K. G., and Dymecki, S. M. (2019). Embracing diversity in the 5-HT neuronal system. Nat Rev Neurosci 20, 397–424. doi: 10.1038/s41583-019-0151-3.

Pollak Dorocic, I., Fürth, D., Xuan, Y., Johansson, Y., Pozzi, L., Silberberg, G., et al. (2014). A Whole-Brain Atlas of Inputs to Serotonergic Neurons of the Dorsal and Median Raphe Nuclei. Neuron 83, 663–678. doi: 10.1016/J.NEURON.2014.07.002.

Ren, J., Friedmann, D., Xiong, J., Liu, C. D., Ferguson, B. R., Weerakkody, T., et al. (2018). Anatomically Defined and Functionally Distinct Dorsal Raphe Serotonin Sub-systems. Cell 175, 472-487.e20. doi: 10.1016/J.CELL.2018.07.043.

Richards, C. D., Shiroyama, T., and Kitai, S. T. (1997). Electrophysiological and immunocytochemical characterization of GABA and dopamine neurons in the substantia nigra of the rat. Neuroscience 80, 545–557. doi: 10.1016/S0306-4522(97)00093-6.

Ritter, J., Lewis, L., Mant, T., and Ferro, A. (2008). A textbook of clinical pharmacology and therapeutics. CRC Press.

Schultz, W., Dayan, P., and Montague, P. R. (1997). A neural substrate of prediction and reward. Science 275, 1593–1599.

Shepard, P. D., and Bunney, B. S. (1991). Repetitive firing properties of putative dopamine-containing neurons in vitro: regulation by an apamin-sensitive Ca2+-activated K+ conductance. Experimental Brain Research 1991 86:1 86, 141–150. doi: 10.1007/BF00231048.

Strogatz, S. H. (2018). Nonlinear dynamics and chaos with student solutions manual: With applications to physics, biology, chemistry, and engineering. CRC press.

Tan, K. R., Yvon, C., Turiault, M., Mirzabekov, J. J., Doehner, J., Labouèbe, G., et al. (2012). GABA Neurons of the VTA Drive Conditioned Place Aversion. Neuron 73, 1173–1183. doi: 10.1016/j.neuron.2012.02.015.

Tian, J., Huang, R., Cohen, J. Y., Osakada, F., Kobak, D., Machens, C. K., et al. (2016). Distributed and mixed information in monosynaptic inputs to dopamine neurons. Neuron 91, 1374–1389.

Tuckwell, H. C., and Penington, N. J. (2014). Computational modeling of spike generation in serotonergic neurons of the dorsal raphe nucleus. Progress in Neurobiology 118, 59–101. doi: 10.1016/j.pneurobio.2014.04.001.

Valencia-Torres, L., Olarte-Sanchez, C. M., Lyons, D. J., Georgescu, T., Greenwald-Yarnell, M., Myers, M. G. J., et al. (2017). Activation of Ventral Tegmental Area 5-HT2C Receptors Reduces Incentive Motivation. Neuropsychopharmacology 42, 1511–1521. doi: 10.1038/npp.2016.264.

Wang, D.-H., and Wong-Lin, K. (2013). Comodulation of dopamine and serotonin on prefrontal cortical rhythms: a theoretical study. Frontiers in Integrative Neuroscience 7, 1–19. doi: 10.3389/fnint.2013.00054.

Wang, H. L., Zhang, S., Qi, J., Wang, H., Cachope, R., Mejias-Aponte, C. A., et al. (2019). Dorsal Raphe Dual Serotonin-Glutamate Neurons Drive Reward by Establishing Excitatory Synapses on VTA Mesoaccumbens Dopamine Neurons. Cell Reports 26, 1128-1142.e7. doi: 10.1016/j.celrep.2019.01.014.

Watabe-Uchida, M., Eshel, N., and Uchida, N. (2017). Neural Circuitry of Reward Prediction Error. Annu Rev Neurosci 40, 373–394. doi: 10.1146/annurev-neuro-072116-031109.

Watabe-Uchida, M., Zhu, L., Ogawa, S. K., Vamanrao, A., and Uchida, N. (2012). Whole-brain mapping of direct inputs to midbrain dopamine neurons. Neuron 74, 858–873.

Weissbourd, B., Ren, J., DeLoach, K. E., Guenthner, C. J., Miyamichi, K., and Luo, L. (2014). Presynaptic partners of dorsal raphe serotonergic and GABAergic neurons. Neuron 83, 645–662.

Whitacre, J. M. (2010). Degeneracy: a link between evolvability, robustness and complexity in biological systems. Theoretical Biology and Medical Modelling 7, 1–17.

Wong-Lin, K., Joshi, A., Prasad, G., and McGinnity, T. M. (2012). Network properties of a computational model of the dorsal raphe nucleus. Neural Networks 32, 15–25. doi: 10.1016/j.neunet.2012.02.009.

Wong-Lin, K., Wang, D.-H., Moustafa, A. A., Cohen, J. Y., and Nakamura, K. (2017). Toward a multiscale modeling framework for understanding serotonergic function. Journal of Psychopharmacology, 026988111769961. doi: 10.1177/0269881117699612.

Xu, P., He, Y., Cao, X., Valencia-Torres, L., Yan, X., Saito, K., et al. (2017). Activation of serotonin 2C receptors in dopamine neurons inhibits binge-like eating in mice. Biol Psychiatry 81, 737–747.

Zhong, W., Li, Y., Feng, Q., and Luo, M. (2017). Learning and stress shape the reward response patterns of serotonin neurons. Journal of Neuroscience 37, 8863–8875.

Zhou, H., Wong-Lin, K., and Wang, D.-H. (2018). Parallel Excitatory and Inhibitory Neural Circuit Pathways Underlie Reward-Based Phasic Neural Responses. Complexity 2018.

Zhou, L., Liu, M. Z., Li, Q., Deng, J., Mu, D., and Sun, Y. G. (2017). Organization of Functional Long-Range Circuits Controlling the Activity of Serotonergic Neurons in the Dorsal Raphe Nucleus. Cell Reports 18, 3018–3032. doi: 10.1016/J.CELREP.2017.02.077.

