## Supplementary Materials for "Degeneracy and stability in neural circuits of dopamine and serotonin neuromodulators: A theoretical consideration"

Behera et al.

### Supplementary Materials

#### *Degeneracy and stability in neural circuits of dopamine and serotonin neuromodulators: A theoretical consideration*

Chandan K. Behera[1,^], Alok Joshi[1], Da-Hui Wang[2,3], Trevor Sharp[4] and KongFatt Wong-Lin[1,*]

[1]*Intelligent Systems Research Centre, School of Computing, Engineering and Intelligent Systems, Ulster University, Derry~Londonderry, Northern Ireland, UK*

[2]*State Key Laboratory of Cognitive Neuroscience and Learning, Beijing Normal University, Beijing, China*

[3]*School of Systems Science, Beijing Normal University, Beijing, China*

[4]*Department of Pharmacology, University of Oxford, Oxford, UK*

*^Current address: Yale School of Medicine, New Haven, Connecticut, USA*

**Abstract**

Degenerate neural circuits perform the same function despite being structurally different. However, it is unclear whether neural circuits with interacting neuromodulator sources can themselves be degenerate while maintaining the same neuromodulatory function. Here, we address this by computationally modelling the neural circuits of neuromodulators serotonin and dopamine, local glutamatergic and GABAergic interneurons, and their possible interactions, under reward/punishment-based conditioning tasks. The neural modelling is constrained by relevant experimental studies of the VTA or DRN system using e.g. electrophysiology, optogenetics, and voltammetry. We first show that a single parsimonious, sparsely connected neural circuit model can recapitulate several separate experimental findings that indicated diverse, heterogeneous, distributed and mixed DRN-VTA neuronal signalling in reward and punishment tasks. The inability for this model to recapitulate all observed neuronal signalling suggests potentially multiple circuits acting in parallel. Then using computational simulations and dynamical systems analysis, we demonstrate that several different stable circuit architectures can produce the same observed network activity profile, hence demonstrating degeneracy. Due to the extensive D2-mediated connections in the investigated circuits, we simulate D2 receptor agonist by increasing the connection strengths emanating from the VTA DA neurons. We found that the simulated D2 agonist can distinguish among sub-groups of the degenerate neural circuits based on substantial deviations in specific neural populations' activities in reward and punishment conditions. This forms a testable model prediction using pharmacological means. Overall, this theoretical work suggests the plausibility of degeneracy within neuromodulator circuitry and has important implication for the stable and robust maintenance of neuromodulatory functions.

In this Supplementary Materials, we provide additional information and data to support the results presented in the main text. The content is organized as follows.

First, we provide in Supplementary Figure 1, a detailed labelled connectivity of the highly connected DRN-VTA circuit model in model architecture a in Fig. 4 of the main text. It also illustrates self-connectivity within each neural population which is not explicitly shown in Figs. 1 and 4 in the main text. Table 1 summarizes the experimental outcomes of the activities of the neuronal populations in DRN-VTA, as illustrated in Figure 1B of main text. Then, Table 2 explicitly shows the range of acceptable values of the internal connectivity strengths for all the degenerate models investigated (Figs. 2 and 4 in main text). This is followed by Table 3 which presents the experimental support for the selected acceptable ranges for the activity profiles of the degenerate neural circuit models (Fig. 4 in main text), and the evaluation of changes in activities due to simulation of pharmacological administration (Fig. 6 in main text). Table 4 is accompanied by a list of additional references (note: some of which are not found in the main text). Table 5 shows the specific degenerate model architectures and #'s mentioned in Figs. 4 and 5 in the main text. The models are based on their distinct connectivity, 5-HT neuronal types, and signs of specific connections. This is then followed by five tables consisting of detailed percentage changes in the models' neural population firing rates due to simulated $D_2$ agonist with increasing dosage level, with Table 5 having the default values. The Supplementary Information ends with a Supplementary Method that provides more detailed mathematical derivation of the steady states of the neural circuit models.

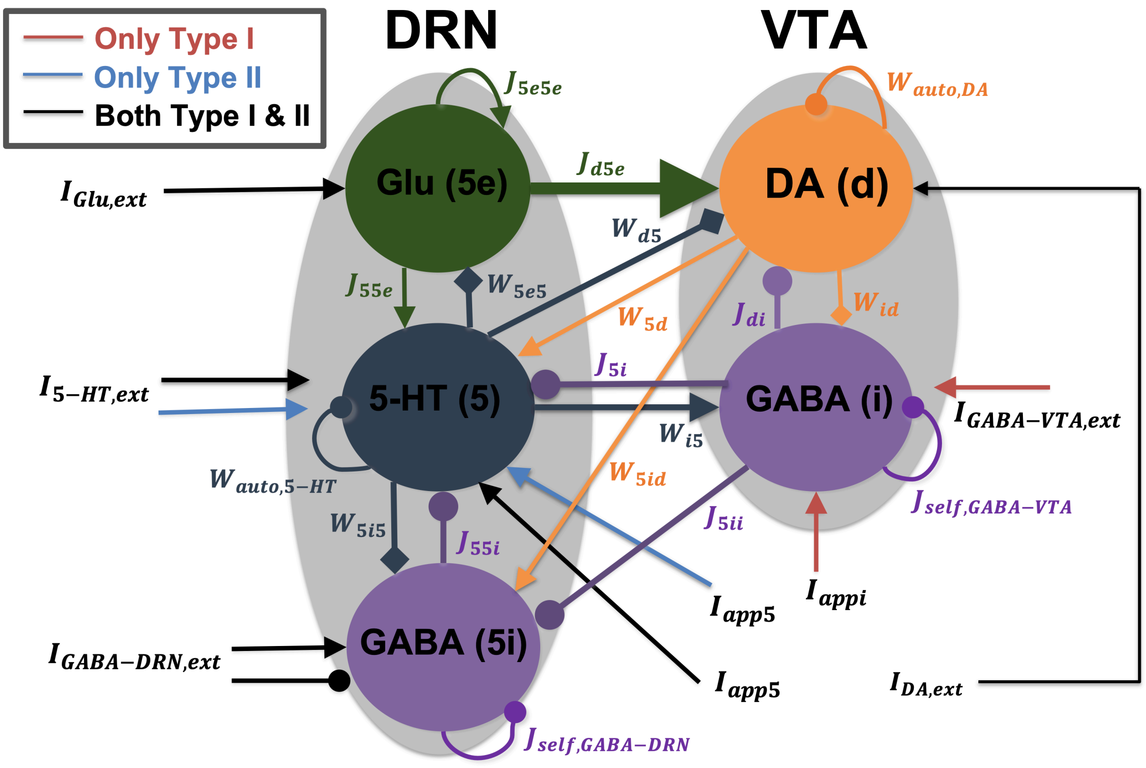


**Supplementary Figure 1. Detailed connectivity and labelling for the highly connected DRN-VTA model with model architecture 'a' in Fig. 4 in main text.** Input currents are labelled with $I$'s, and relative weights of connectivity with $W$'s and $J$'s, $W$'s if 5-HT/DA-mediated connections associated; $J$'s if glutamatergic/GABA-mediated connections. These define the subscripts described in the equations in Materials and Methods. Colours of connectivity are based on sources of the connectivity. Thicker arrows (not scaled) denote stronger connectivity; the range of values of the relative weights are found in Supplementary Information, Table 2. Ionotropic receptor mediated self-connection strengths within DRN Glu, DRN GABA and VTA GABA neurons are fixed for all simulations; however, in drug simulations, DA neuronal self-connection (auto-inhibition) is increased (see main text).

| Neuronal populations | Firing activities based on Type-1 5-HT neurons | Firing activities based on Type-II 5-HT neurons |
|---|---|---|
| 5-HT | Phasic increase of firing rates for both reward during cue. No change in baseline activities for punishment between cue and outcome (Cohen et al.,2015, Li et al., 2016, Liu et al. 2014) | Phasic increase in reward and punishment during cue and outcome respectively. The baseline activity during reward remains the same between cue and outcome (Cohen et al.,2015, Li et al., 2016, Liu et al. 2014) |
| DA | Phasic decrease during outcome for punishment (Tan et al., 2012). Phasic increase during cue for reward (Cohen et al., 2012) | No change in the baseline activities between reward and punishment (Cohen et al.,2015) |
| DRN-GABA | Phasic increase during outcome for punishment (Li et al., 2016) | Phasic increase and decrease during outcomes for punishment and reward respectively (Li et al., 2016) |
| DRN Glu | Same as DA activities in cue during reward (McDevitt et al., 2014) | Same as DA activities in cue during reward (McDevitt et al., 2014) |
| VTA-GABA | Phasic rise at the outcome during punishment. Exponential rise from cue to outcome during reward (Tan et al., 2012) | Activities increase from cue to outcome for reward and punishment (Cohen et al, 2012) |

**Table 1. Experimental observation on across trial reward/punishment based conditioning tasks.** The table summarizes the experimental outcomes of the activities of the neuronal populations in DRN-VTA, as illustrated in Figure 1B of main text.

| | a | b | c | d | e | f | g | h | i | j | k | l* |
|---|---|---|---|---|---|---|---|---|---|---|---|---|
| $J_{55e}$ | 10 | 6 | 5 | 5 | 5 | 5 | 5 | 5 | 5 | 5 | 5 | 5 |
| $J_{5i}$ | 0-1 | 0 | 0 | 0 | 0 | 0 | 0 | 0 | 0 | 0 | 0 | 0 |
| $J_{55i}$ | 0-0.1 | 0-0.1 | 0-0.1 | 0-0.1 | 0-0.1 | 0-0.1 | 0 | 0 | 0 | 0 | 0 | 0.1 |
| $W_{5d}$ | 0-30 | 0-10 | 0-10 | 0-10 | 0-10 | 0 | 0 | 0-10 | 0 | 0 | 0 | 0-10 |
| $J_{d5e}$ | 100 | 100 | 100 | 100 | 100 | 100 | 100 | 100 | 100 | 100 | 100 | 100 |
| $J_{di}$ | 25 | 25 | 25 | 25 | 25 | 25 | 25 | 25 | 25 | 25 | 25 | 25 |
| $W_{d5}$ | 0-1 | 0-1 | 0-1 | 0-1 | 0 | 0 | 0 | 0 | 0-2 | 0-2 | 0 | 0-1 |
| $W_{5e5}$ | 0-0.1 | 0-0.1 5HT to Glu (ex): 0-1 | 0 | 0 | 0 | 0 | 0 | 0 | 0 | 0 | 0 | 0-0.1 |
| $W_{i5}$ | 0-20 | 0-20 | 0-20 | 0-20 | 0-20 | 0-20 | 0-20 | 0-20 | 0-20 | 0-20 | 0-20 | 0-20 |
| $W_{id}$ | 0-2 | 0-2 | 0-2 | 0 | 0 | 0 | 0 | 0 | 0 | 0 | 0 | 0-2 |
| $W_{5i5}$ | p: 0-0.2 r:0-4 | 0-4 | 0-4 | 0-10 | 0-10 | 0-10 | 0-10 | 0-10 | 0-10 | 0-10 | 0-10 | p: 0-4 r:0-3 |
| $W_{5id}$ | p:0-10 r:0-20 | 0-10 | 0-10 | 0-10 | 0 | 0 | 0-20 | 0 | 0 | 0 | 0 | 0-10 |
| $J_{self,Glu}$ | 0.5 | 0.5 | 0.5 | 0.5 | 0.5 | 0.5 | 0.5 | 0.5 | 0.5 | 0.5 | 0.5 | 0.5 |
| $J_{self,GABA-DRN}$ | 0.5 | 0.5 | 0.5 | 0.5 | 0.5 | 0.5 | 0.5 | 0.5 | 0.5 | 0.5 | 0.5 | 0.5 |
| $W_{5ii}$ | p:0.4 r:0.1 | 0 | 0 | 0 | 0 | 0 | 0 | 0 | 0 | 0 | 0 | 0.4 |
| $J_{self,GABA-VTA}$ | 10 | 10 | 10 | 10 | 10 | 10 | 10 | 10 | 10 | 10 | 10 | 10 |
| $W_{auto,DA}$ | 1 | 1 | 1 | 1 | 1 | 1 | 1 | 1 | 1 | 1 | 1 | 1 |
| $W_{auto,5-HT}$ | 1 | 1 | 1 | 1 | 1 | 1 | 1 | 1 | 1 | 1 | 1 | 1 |

**Table 2. Summary of model parameter values of internal connections considered for the degenerate DRN-VTA circuit models (Fig. 4 in main text).** Column: Model architectures; row: connection strength (see Materials and methods in main text for definitions; and Supplementary Information, Fig. 1). These values reflect either single values used or a range of values such that their corresponding neural activity profiles fit within the degeneracy criteria (Materials and methods). These values are used with Type I or II 5-HT neuronal model, punishment or reward unless specified with p or r, respectively, and excitatory or inhibitory connections unless specified with + or −, respectively. Self-connections are not shown as they are constants except for the DA neurons during $D_2$ agonist drug simulation cases (see main text). See Supplementary Information, Table 4 for detailed division and grouping of model #'s.

| ***Brain region & neuron type*** | ***Firing rate*** | ***Reference*** |
|---|---|---|
| **VTA DA** | ~ 4 - 6 Hz | *Cheer, J. F., Kendall, D. A., Mason, R. & Marsden, C. A. Differential cannabinoid-induced electrophysiological effects in rat ventral tegmentum. *Neuropharmacology* **44**: 633–641 (2003);<br><br>[+]Ungless, M. A., Magill, P. J., Bolam, J. P. Uniform inhibition of dopamine neurons in the ventral tegmental area by aversive stimuli. *Science* **303**(5666): 2040-2042 (2004);<br><br>*Li, W., Doyon, W. M. & Dani, J. A. Quantitative unit classification of ventral tegmental area neurons in vivo. *J. Neurophysiol*. **107**(10):2808–2820 (2012);<br><br>[+]Tolu, S., Eddine, R., Marti, F., David, V., Graupner, M., Pons, S., Baudonnat, M., Husson, M., Besson, M., Reperant, C., Zemdegs, J., Pages, C., Hay, Y. A. H., Lobolez, B., Caboche, J., Gutkin, B., Gardier, A. M., Changeaux, J.-P., Faure, P. & Maskos, U. Co-activation of VTA DA and GABA neurons mediates nicotine reinforcement. *Mol. Psychiatry* **18**(3):382-392 (2013);<br><br>[+]Tian, J., Huang, R., Cohen, J. Y., Osakada, F., Kobak, D., Machens, C. K., Callaway, E. M., Uchida, N. & Watabe-Uchida, M. Distributed and mixed information in monosynaptic inputs to dopamine neurons. *Neuron* **91**:1374-1389 (2016);<br><br>[+]Stauffer, W. R., Lak, A., Yang, A., Borel, M., Paulsen, O., Boyden, E. S. & Schultz, W. Dopamine neuron-specific optogenetic stimulation in rhesus macaques. *Cell* **166**:1564-1571 (2016). |
| **VTA GABA** | ~ 12 - 17 Hz | [+]Steffensen, S. C., Svingos, A. L., Pickel, V. M. & Henriksen, S. J. Electrophysiological characterization of GABAergic neurons in the ventral tegmental area. *J. Neurosci.* **18**(19):8003-8015 (1998);<br><br>[+]Steffensen, S. C., Taylor, S. R., Horton, M. L., Barber, E. N., Lyle, L. T., Stobbs, S. H. & Allsion, D. W. (Cocaine disinhibits dopamine neurons in the ventral tegmental area via use-dependent blockade of GABA neuron voltage-sensitive sodium channels. *Eur. J. Neurosci.* **28**(10):2028-2040 (2008);<br><br>[+]Steffensen, S. C., Walton, C. H., Hansen, D. M., Yorgason, J. T., Gallegos, R. A. & Criado, J. R. Contingent and non-contingent effects of low-dose ethanol on GABA neuron activity in the ventral tegmental area. *Pharmacol. Biochem. Behav*. **92**(1):68-75 (2009);<br><br>[+]Cohen, J., Haesler, S., Vong, L., Lowell, B. B. & Uchida, N. Neuron-type specific signals for reward and punishment in the ventral tegmental area. *Nature* **482**(7383):85-88 (2012);<br><br>[+] Tolu, S., Eddine, R., Marti, F., David, V., Graupner, M., Pons, S., Baudonnat, M., Husson, M., Besson, M., Reperant, C., Zemdegs, J., Pages, C., Hay, Y. A. H., Lobolez, B., Caboche, J., Gutkin, B., Gardier, A. M., Changeaux, J.-P., Faure, P. & Maskos, U. Co-activation of VTA DA and GABA neurons mediates nicotine reinforcement. *Mol. Psychiatry* **18**(3):382-392 (2013). |
| **DRN 5-HT** | ~ 3 - 5 Hz | [+]Allers, K. A. & Sharp, T. Neurochemical and anatomical identification of fast- and slow-firing neurones in the rat dorsal raphe nucleus using juxtacellular labelling methods in vivo. *Neuroscience* **122**(1):193–204 (2003);<br><br>*Judge, S. J. & Gartside, S. E. Firing of 5-HT neurones in the dorsal and median raphe nucleus in vitro shows differential alpha1-adrenoreceptor and 5-HT1A receptor modulation. *Neurochem. Int.* **48**(2):100-107 (2006);<br><br>[+]Hajós, M., Allers, K. A., Jennings, K., Sharp, T., Charette, G., Sík, A. & Kocsis, B. Neurochemical identification of stereotypic burst-firing neurons in the rat dorsal raphe nucleus using juxtacellular labelling methods. *Eur. J. Neurosci*. **25**(1):119–126 (2007);<br><br>[+]Ranade, S. P. & Mainen, Z. F. Transient firing of dorsal raphe neurons encodes diverse and specific sensory, motor, and reward events. *J. Neurophysiol.* **102**(5):3026-3037 (2009);<br><br>[+]Cohen, J. Y., Amoroso, M. W. & Uchida, N. Serotonergic neurons signal reward and punishment on multiple timescales. *eLife* **4**:e06346 (2015). doi:10.7554/eLife.06346;<br><br>[+]Li, Y., Zhong, W., Wang, D., Feng, Q., Liu, Z., Zhou, J., Jia, C., Hu, F., Zeng, J., Guo, Q., |

| | | |
|---|---|---|
| | | Fu, L. & Luo, M. Serotonin neurons in the dorsal raphe nucleus encode reward signals. *Nat. Commun.* **7**:10503 (2016). https://doi.org/10.1038/ncomms10503;<br><br>*Mlinar, B., Montalbano, A., Piszczek, L., Gross, C. & Corradetti, R. Firing properties of genetically identified dorsal raphe serotonergic neurons in brain slices. *Front. Cell. Neurosci.* **10**:195 (2016). doi: 10.3389/fncel.2016.00195;<br><br>*Srejic, L. R., Wood, K. M., Zeqja, A., Hashemi, P., Hutchison, W. D. Modulation of serotonin dynamics in the dorsal raphe nucleus via high frequency medial prefrontal cortex stimulation. *Neurobiol. Dis.* **94**:129-138 (2016). |
| **DRN GABA** | ~ 15 - 25 Hz | +Allers, K. A. & Sharp, T. Neurochemical and anatomical identification of fast- and slow-firing neurones in the rat dorsal raphe nucleus using juxtacellular labelling methods in vivo. *Neuroscience* **122**(1):193–204 (2003);<br><br>+Sakai K. Sleep-waking discharge profiles of dorsal raphe nucleus neurons in mice. *Neuroscience* **197**:200-224 (2011);<br><br>+Challis, C., Boulden, J., Veerakumar, A., Espallergues, J., Vassoler, F. M., Pierce, R. C., Beck, S. G. & Berton, O. Raphe GABAergic neurons mediate the acquisition of avoidance after social defeat. *J. Neurosci*. **33**(35):13978-13988a (2013);<br><br>+Li, Y., Zhong, W., Wang, D., Feng, Q., Liu, Z., Zhou, J., Jia, C., Hu, F., Zeng, J., Guo, Q., Fu, L. & Luo, M. Serotonin neurons in the dorsal raphe nucleus encode reward signals. *Nat. Commun.* **7**:10503 (2016). https://doi.org/10.1038/ncomms10503;<br><br>*Hernández-Vázquez, F., Garduñom J. & Hernández-López, S. GABAergic modulation of serotonergic neurons in the dorsal raphe nucleus. *Rev. Neurosci.* **30**(3):289-303 (2018). |
| **DRN Glu** | ~ 3 - 5 Hz | +Taylor, N. E., Pei, J., Zhang, Vlasov, K. Y., Davis, T., Taylor, E., Weng, F.-J., Van Dort, C. J., Solt, K. & Brown, E. N. The role of glutamatergic and dopaminergic neurons in the periaqueductal gray/dorsal raphe: separating analgesia and anxiety. *eNeuro* **6**(1):ENEURO.0018-18.2019 (2019). doi:10.1523/ENEURO.0018-18.2019. |

**Table 3.** Firing rate ranges estimated based on relevant references. *: *in vitro* recording data, +: *in vivo* recording data. Only baseline firing rates were deduced due to the higher variation in stimulus-evoked activities. Note: references shown are samples but do not reflect the complete list in the literature. These ranges provide a leverage to identify acceptable activity profile deviations for the degenerate neural circuit models in Fig. 4 and the effects of drugs in Fig. 6 in main text.

| Model Architecture (Fig.4) | | | | Type-I 5-HT neurons | | Type-II 5-HT neurons | |
|---|---|---|---|---|---|---|---|
| | | | | Punishment | Reward | Punishment | Reward |
| | | | | Model # | | | |
| ***A*** | DA→VTA GABA (-)<br>5HT→DRN GABA (-) | 5HT→DA (-) | 5HT→Glu (-) | 1 | 2 | 3 | 4 |
| | | | 5HT→Glu (+) | 5 | 6 | 7 | 8 |
| | | 5HT→DA (+) | 5HT→Glu (-) | 9 | 10 | 11 | 12 |
| | | | 5HT→Glu (+) | 13 | 14 | 15 | 16 |
| | DA→VTA GABA (+)<br>5HT→DRN GABA (+) | 5HT→DA (-) | 5HT→Glu (-) | 17 | 18 | 19 | 20 |
| | | | 5HT→Glu (+) | 21 | 22 | 23 | 24 |
| | | 5HT→DA (+) | 5HT→Glu (-) | 25 | 26 | 27 | 28 |
| | | | 5HT→Glu (+) | 29 | 30 | 31 | 32 |
| ***B*** | 5HT→Glu (-) | | | 33 | 34 | 35 | 36 |
| | 5HT→Glu (+) | | | 37 | 38 | 39 | 40 |
| ***C*** | - | | | 41 | 42 | 43 | 44 |
| ***D*** | - | | | 45 | 46 | 47 | 48 |
| ***E*** | - | | | 49 | 50 | 51 | 52 |
| ***F*** | - | | | 53 | 54 | 55 | 56 |
| ***G*** | - | | | 57 | 58 | 59 | 60 |
| ***H*** | - | | | 61 | 62 | 63 | 64 |
| ***I*** | - | | | 65 | 66 | 67 | 68 |
| ***J*** | - | | | 69 | 70 | 71 | 72 |
| **K** | - | | | 73 | 74 | 75 | 76 |
| **L** | 5HT→DA (-) | | | 77 | 78 | 79 | 80 |
| | 5HT→DA (+) | | | 81 | 82 | 83 | 84 |

**Table 4. Degenerate DRN-VTA model #'s and architectures based on specific connectivity and 5-HT neuronal types and reward/punishment task.** The model architectures and #'s (right side of table) discussed in Figs. 4 and 5 in main text are dependent on specific connectivity type (left side of table), 5-HT neuronal type and punishment/reward task (right side of table). Arrows denote connections. Sub-divisions or branching of the connectivity types from the model architectures are also indicated. + / – : effectively positive or negative connection. Some boxes are not filled due to those connections being absent in those models (e.g. model architecture 'k' has no 5-HT→Glu and 5-HT→DA connections; see Fig. 4 in main text).

| Arc | | | | Type-I 5-HT neurons | | | | | | | | | | Type-II 5-HT neurons | | | | | | | | | |
|---|---|---|---|---|---|---|---|---|---|---|---|---|---|---|---|---|---|---|---|---|---|---|---|
| a | Connections/Type | | | Punishment | | | | | Reward | | | | | Punishment | | | | | Reward | | | | |
| | | | | DA VTA | 5-HT DRN | GA DRN | GA VTA | Glu VTA | DA VTA | 5-HT DRN | GA DRN | GA VTA | Glu VTA | DA VTA | 5-HT DRN | GA DRN | GA VTA | Glu VTA | DA VTA | 5-HT DRN | GA DRN | GA VTA | Glu VTA |
| | DA→ GA-VTA (-) | 5-HT→ DA (-) | 5-HT→ Glu (-) | 4.03 | 3.07 | 5.60 | 0.93 | 0.36 | 7.27 | 3.70 | 9.76 | 2.51 | 0.60 | 2.36 | 3.30 | 6.08 | 1.00 | 0.38 | 5.23 | 3.38 | 13.4 | 2.97 | 0.70 |
| | | | 5-HT→ Glu (+) | 1.64 | 2.82 | 5.58 | 0.86 | 0.36 | 3.15 | 4.77 | 9.87 | 1.70 | 0.50 | 0.41 | 3.04 | 6.06 | 0.92 | 0.38 | 1.77 | 3.88 | 13.42 | 1.50 | 0.59 |
| | 5-HT→ GA-VTA (-) | 5-HT→ DA (+) | 5-HT→ Glu (-) | 2.02 | 3.10 | 5.60 | 0.94 | 0.36 | 1.44 | 3.33 | 6.08 | 1.01 | 0.38 | 0.73 | 3.33 | 6.08 | 1.01 | 0.38 | 1.27 | 4.12 | 13.42 | 1.67 | 0.59 |
| | | | 5-HT→ Glu (+) | 1.64 | 2.82 | 5.58 | 0.86 | 0.36 | 6.71 | 4.48 | 10.04 | 2.47 | 0.48 | 0.41 | 3.04 | 6.06 | 0.92 | 0.38 | 4.24 | 3.64 | 13.45 | 2.53 | 0.57 |
| | DA→ GA-VTA (+) | 5-HT→ DA (-) | 5-HT→ Glu (-) | 3.98 | 3.24 | 13.17 | 0.93 | 0.36 | 7.46 | 3.68 | 10.51 | 2.55 | 0.60 | 2.35 | 3.47 | 14.24 | 0.99 | 0.38 | 5.35 | 3.36 | 13.57 | 3.02 | 0.70 |
| | | | 5-HT→ Glu (+) | 1.63 | 2.99 | 13.18 | 0.85 | 0.36 | 2.95 | 4.79 | 10.50 | 1.65 | 0.50 | 0.41 | 3.21 | 14.25 | 0.92 | 0.38 | 1.72 | 3.89 | 13.55 | 1.45 | 0.59 |
| | 5-HT→ GA-VTA (+) | 5-HT→ DA (+) | 5-HT→ Glu (-) | 2.02 | 3.27 | 13.16 | 0.93 | 0.36 | 1.44 | 5.02 | 10.49 | 1.79 | 0.49 | 0.73 | 3.50 | 14.24 | 1.00 | 0.38 | 1.31 | 4.13 | 13.55 | 1.62 | 0.59 |
| | | | 5-HT→ Glu (+) | 1.63 | 2.99 | 13.18 | 0.85 | 0.36 | 6.49 | 4.49 | 10.35 | 2.42 | 0.48 | 0.41 | 3.21 | 14.25 | 0.92 | 0.38 | 4.11 | 3.65 | 13.52 | 2.48 | 0.57 |
| b | 5-HT→Glu (-) | | | 2.86 | 1.12 | 6.01 | 0.40 | 0.38 | 5.25 | 0.98 | 10.24 | 0.60 | 0.55 | 1.59 | 1.11 | 6.48 | 0.41 | 0.40 | 4.07 | 0.86 | 12.94 | 0.70 | 0.64 |
| | 5-HT→Glu (+) | | | 9.89 | 0.20 | 6.08 | 0.06 | 3.75 | 12.68 | 0.53 | 10.37 | 0.28 | 5.41 | 4.93 | 0.23 | 6.56 | 0.06 | 3.98 | 10.2 | 0.59 | 13.08 | 0.36 | 6.36 |
| c | - | | | 2.59 | 0.66 | 6.12 | 0.18 | 0.00 | 3.87 | 0.26 | 10.48 | 0.17 | 0.00 | 1.33 | 0.59 | 6.60 | 0.19 | 0.00 | 3.24 | 0.23 | 13.22 | 0.16 | 0.00 |
| d | - | | | 3.12 | 0.50 | 0.06 | 0.16 | 0.00 | 5.36 | 0.43 | 0.05 | 0.23 | 0.00 | 1.88 | 0.48 | 0.05 | 0.16 | 0.00 | 4.36 | 0.38 | 0.08 | 0.27 | 0.00 |
| e | - | | | 1.50 | 0.52 | 0.08 | 0.17 | 0.00 | 0.35 | 0.18 | 0.08 | 0.12 | 0.00 | 0.25 | 0.45 | 0.09 | 0.18 | 0.00 | 0.17 | 0.15 | 0.07 | 0.10 | 0.00 |
| f | - | | | 1.75 | 1.03 | 0.16 | 0.32 | 0.00 | 1.25 | 0.57 | 0.26 | 0.35 | 0.00 | 0.49 | 0.94 | 0.17 | 0.34 | 0.00 | 0.92 | 0.46 | 0.30 | 0.38 | 0.00 |
| g | - | | | 1.23 | 0.00 | 0.27 | 0.00 | 0.00 | 0.00 | 0.00 | 0.43 | 0.00 | 0.00 | 0.00 | 0.00 | 0.28 | 0.00 | 0.00 | 0.00 | 0.00 | 0.49 | 0.00 | 0.00 |
| h | - | | | 1.49 | 0.50 | 0.08 | 0.16 | 0.00 | 0.91 | 0.44 | 0.17 | 0.23 | 0.00 | 0.24 | 0.49 | 0.08 | 0.17 | 0.00 | 0.75 | 0.39 | 0.23 | 0.28 | 0.00 |
| i | - | | | 4.56 | 0.00 | 0.00 | 0.00 | 0.00 | 8.85 | 0.00 | 0.00 | 0.00 | 0.00 | 3.25 | 0.00 | 0.00 | 0.00 | 0.00 | 7.18 | 0.00 | 0.00 | 0.00 | 0.00 |
| J | - | | | 4.43 | 0.00 | 0.00 | 0.00 | 0.00 | 8.85 | 0.00 | 0.00 | 0.00 | 0.00 | 3.25 | 0.00 | 0.00 | 0.00 | 0.00 | 7.19 | 0.00 | 0.00 | 0.00 | 0.00 |
| k | - | | | 1.23 | 0.00 | 0.00 | 0.00 | 0.00 | 0.00 | 0.00 | 0.00 | 0.00 | 0.00 | 0.00 | 0.00 | 0.00 | 0.00 | 0.00 | 0.00 | 0.00 | 0.00 | 0.00 | 0.00 |
| l | 5-HT→DA (-) | | | 1.44 | 0.72 | 5.44 | 0.14 | 0.37 | 4.31 | 0.26 | 4.08 | 0.25 | 0.53 | 1.56 | 0.66 | 5.91 | 0.19 | 0.39 | 3.75 | 0.29 | 12.57 | 0.39 | 0.63 |
| | 5-HT→DA (+) | | | 1.53 | 0.71 | 5.45 | 0.15 | 1.09 | 1.09 | 4.09 | 0.25 | 0.53 | 1.09 | 0.27 | 0.65 | 5.91 | 0.19 | 0.39 | 1.17 | 0.29 | 12.56 | 0.39 | 0.63 |

**Table 5. Percentage changes in neural population firing rates with $D_2$ receptor-mediated connection strengths changed by a factor of 1 (X=1; default condition).** Time averaged percentage changes in neural population activities. Arc: Architecture type. Nomenclatures same as in Supplementary Information, Table 4, except GA, which is an abbreviation of GABA.

| Arc | Type-I 5-HT neurons | Type-II 5-HT neurons |
|---|---|---|

| a | Connections/Type | | | Punishment | | | | | Reward | | | | | Punishment | | | | | Reward | | | | |
|---|---|---|---|---|---|---|---|---|---|---|---|---|---|---|---|---|---|---|---|---|---|---|---|
| | | | | DA VTA | 5HT DRN | GA DRN | GA VTA | Glu VTA | DA VTA | 5HT DRN | GA DRN | GA VTA | Glu VTA | DA VTA | 5HT DRN | GA DRN | GA VTA | Glu VTA | DA VTA | 5HT DRN | GA DRN | GA VTA | Glu VTA |
| | **DA→ GA-VTA (-)** | **5-HT→ DA (-)** | **5-HT→ Glu (-)** | 36.55 | 2.75 | 6.05 | 0.73 | 0.39 | 43.42 | 8.21 | 11.98 | 5.23 | 0.66 | 37.31 | 2.26 | 6.52 | 0.74 | 0.41 | 43.36 | 7.21 | 13.76 | 5.92 | 0.77 |
| | | | **5-HT→ Glu (+)** | 38.16 | 2.95 | 6.03 | 0.80 | 0.39 | 40.60 | 1.92 | 12.17 | 0.46 | 0.55 | 39.23 | 2.47 | 6.49 | 0.81 | 0.42 | 41.79 | 1.73 | 13.80 | 1.04 | 0.66 |
| | **5-HT→ GA-VTA (-)** | **5-HT→ DA (+)** | **5-HT→ Glu (-)** | 39.09 | 2.61 | 6.04 | 0.69 | 0.39 | 41.50 | 2.12 | 6.51 | 0.69 | 0.41 | 40.17 | 2.12 | 6.51 | 0.69 | 0.41 | 42.29 | 1.67 | 13.79 | 0.82 | 0.65 |
| | | | **5-HT→ Glu (+)** | 38.16 | 2.95 | 6.03 | 0.80 | 0.39 | 41.43 | 2.43 | 12.32 | 0.99 | 0.56 | 39.23 | 2.47 | 6.49 | 0.81 | 0.42 | 42.06 | 2.17 | 13.84 | 1.50 | 0.67 |
| | **DA→ GA-VTA (+)** | **5-HT→ DA (-)** | **5-HT→ Glu (-)** | 36.90 | 2.46 | 14.33 | 1.03 | 0.39 | 43.58 | 8.16 | 12.81 | 5.55 | 0.66 | 37.69 | 1.98 | 15.43 | 1.04 | 0.41 | 43.52 | 7.17 | 14.00 | 6.26 | 0.77 |
| | | | **5-HT→ Glu (+)** | 38.50 | 2.66 | 14.33 | 1.10 | 0.39 | 40.89 | 1.89 | 12.86 | 0.76 | 0.55 | 39.58 | 2.19 | 15.43 | 1.10 | 0.41 | 41.95 | 1.70 | 14.00 | 1.36 | 0.66 |
| | **5-HT→ GA-VTA (+)** | **5-HT→ DA (+)** | **5-HT→ Glu (-)** | 39.42 | 2.32 | 14.30 | 0.98 | 0.39 | 41.79 | 1.78 | 12.82 | 0.58 | 0.55 | 40.47 | 1.84 | 15.40 | 0.99 | 0.41 | 42.44 | 1.66 | 13.99 | 1.14 | 0.65 |
| | | | **5-HT→ Glu (+)** | 38.50 | 2.66 | 14.33 | 1.10 | 0.39 | 41.74 | 2.35 | 12.67 | 1.31 | 0.56 | 39.58 | 2.19 | 15.43 | 1.10 | 0.41 | 42.22 | 2.11 | 13.94 | 1.83 | 0.67 |
| b | **5-HT→Glu (-)** | | | 39.10 | 4.18 | 6.64 | 1.58 | 0.40 | 42.67 | 3.38 | 10.99 | 2.12 | 0.58 | 40.17 | 4.05 | 7.10 | 1.63 | 0.42 | 42.76 | 2.90 | 13.70 | 2.40 | 0.68 |
| | **5-HT→Glu (+)** | | | 33.93 | 3.37 | 6.74 | 1.28 | 3.95 | 37.17 | 3.13 | 11.16 | 1.89 | 5.76 | 34.21 | 3.33 | 7.21 | 1.33 | 4.19 | 39.47 | 2.80 | 13.87 | 2.19 | 6.76 |
| c | - | | | 38.88 | 2.42 | 6.76 | 0.97 | 0.00 | 42.34 | 2.16 | 11.24 | 1.31 | 0.00 | 39.97 | 2.36 | 7.23 | 1.00 | 0.00 | 42.61 | 1.85 | 13.99 | 1.51 | 0.00 |
| d | - | | | 39.13 | 3.58 | 0.39 | 1.16 | 0.00 | 42.68 | 2.85 | 0.23 | 1.58 | 0.00 | 40.20 | 3.44 | 0.36 | 1.21 | 0.00 | 42.80 | 2.44 | 0.38 | 1.81 | 0.00 |
| e | - | | | 37.44 | 2.62 | 0.41 | 0.84 | 0.00 | 40.89 | 2.36 | 0.94 | 1.25 | 0.00 | 38.41 | 2.57 | 0.44 | 0.87 | 0.00 | 41.84 | 2.04 | 1.20 | 1.46 | 0.00 |
| f | - | | | 36.10 | 1.03 | 0.16 | 0.32 | 0.00 | 39.15 | 0.57 | 0.26 | 0.35 | 0.00 | 36.83 | 0.94 | 0.17 | 0.34 | 0.00 | 41.04 | 0.46 | 0.30 | 0.38 | 0.00 |
| g | - | | | 36.48 | 0.00 | 1.96 | 0.00 | 0.00 | 39.53 | 0.00 | 2.82 | 0.00 | 0.00 | 37.27 | 0.00 | 1.98 | 0.00 | 0.00 | 41.25 | 0.00 | 3.09 | 0.00 | 0.00 |
| h | - | | | 37.83 | 3.62 | 0.58 | 1.18 | 0.00 | 41.28 | 2.90 | 1.21 | 1.61 | 0.00 | 38.87 | 3.48 | 0.62 | 1.23 | 0.00 | 42.05 | 2.48 | 1.50 | 1.84 | 0.00 |
| i | - | | | 34.00 | 0.00 | 0.00 | 0.00 | 0.00 | 36.88 | 0.00 | 0.00 | 0.00 | 0.00 | 34.32 | 0.00 | 0.00 | 0.00 | 0.00 | 39.31 | 0.00 | 0.00 | 0.00 | 0.00 |
| j | - | | | 38.95 | 0.00 | 0.00 | 0.00 | 0.00 | 42.20 | 0.00 | 0.00 | 0.00 | 0.00 | 40.05 | 0.00 | 0.00 | 0.00 | 0.00 | 42.69 | 0.00 | 0.00 | 0.00 | 0.00 |
| k | - | | | 36.48 | 0.00 | 0.00 | 0.00 | 0.00 | 39.53 | 0.00 | 0.00 | 0.00 | 0.00 | 37.27 | 0.00 | 0.00 | 0.00 | 0.00 | 41.25 | 0.00 | 0.00 | 0.00 | 0.00 |
| l | **5-HT→DA (-)** | | | 38.81 | 2.36 | 6.06 | 1.15 | 0.39 | 42.36 | 2.16 | 4.91 | 1.59 | 0.56 | 40.05 | 2.30 | 6.53 | 1.24 | 0.41 | 42.65 | 1.93 | 12.49 | 1.92 | 0.66 |
| | **5-HT→DA (+)** | | | 37.85 | 2.39 | 6.07 | 1.16 | 0.39 | 41.33 | 2.20 | 4.93 | 1.61 | 0.56 | 38.89 | 2.33 | 6.53 | 1.25 | 0.41 | 42.08 | 1.96 | 12.48 | 1.95 | 0.66 |

**Table 6. Percentage changes in neural population firing rates with $D_2$ receptor-mediated connection strengths changed by a factor of 10 (X=10).** Time-averaged percentage changes in neural population activities. Arc: Architecture type. Nomenclatures same as in Supplementary Information, Table 4, except GA, which is an abbreviation of GABA. Values in red: Percentage changes beyond the prescribed allowed ranges set by the inclusion criterion.

| Arc | | | | Type-I 5-HT neurons | | | | | | | | | | Type-II 5-HT neurons | | | | | | | | | |
|---|---|---|---|---|---|---|---|---|---|---|---|---|---|---|---|---|---|---|---|---|---|---|---|
| a | Connections/Type | | | **Punishment** | | | | | **Reward** | | | | | **Punishment** | | | | | **Reward** | | | | |
| | | | | **DA VTA** | **5HT DRN** | **GA DRN** | **GA VTA** | **Glu VTA** | **DA VTA** | **5HT DRN** | **GA DRN** | **GA VTA** | **Glu VTA** | **DA VTA** | **5HT DRN** | **GA DRN** | **GA VTA** | **Glu VTA** | **DA VTA** | **5HT DRN** | **GA DRN** | **GA VTA** | **Glu VTA** |
| | **DA→ GA-VTA (-)** | **5-HT→ DA (-)** | **5-HT→ Glu (-)** | 46.15 | 20.25 | 7.22 | 7.07 | 0.52 | 45.66 | 19.37 | 17.63 | 13.97 | 0.88 | 46.15 | 19.04 | 7.65 | 7.28 | 0.55 | 45.63 | 17.02 | 15.97 | 14.61 | 0.98 |
| | | | **5-HT→ Glu (+)** | 46.15 | 20.62 | 7.19 | 7.24 | 0.52 | 45.61 | 11.02 | 17.80 | 7.29 | 0.72 | 46.15 | 19.42 | 7.62 | 7.46 | 0.55 | 45.59 | 9.82 | 16.03 | 8.31 | 0.84 |
| | **5-HT→ GA-VTA (-)** | **5-HT→ DA (+)** | **5-HT→ Glu (-)** | 46.15 | 20.25 | 7.22 | 7.07 | 0.52 | 45.63 | 10.68 | 17.80 | 7.03 | 0.71 | 46.15 | 19.04 | 7.65 | 7.28 | 0.55 | 45.60 | 9.48 | 16.03 | 8.03 | 0.83 |
| | | | **5-HT→ Glu (+)** | 46.15 | 20.62 | 7.19 | 7.24 | 0.52 | 45.68 | 18.59 | 17.91 | 13.01 | 0.85 | 46.15 | 19.42 | 7.62 | 7.46 | 0.55 | 45.65 | 16.35 | 16.09 | 13.77 | 0.96 |
| | **DA→ GA-VTA (+)** | **5-HT→ DA (-)** | **5-HT→ Glu (-)** | 46.15 | 19.28 | 18.73 | 15.04 | 0.88 | 45.67 | 19.28 | 18.73 | 15.04 | 0.88 | 46.15 | 18.49 | 19.36 | 8.48 | 0.54 | 45.64 | 16.94 | 16.45 | 15.72 | 0.98 |
| | | | **5-HT→ Glu (+)** | 46.15 | 20.06 | 18.18 | 8.44 | 0.52 | 45.62 | 10.77 | 18.70 | 8.25 | 0.72 | 46.15 | 18.87 | 19.39 | 8.65 | 0.55 | 45.60 | 9.61 | 16.43 | 9.29 | 0.83 |
| | **5-HT→ GA-VTA (+)** | **5-HT→ DA (+)** | **5-HT→ Glu (-)** | 46.15 | 19.69 | 18.15 | 8.27 | 0.52 | 45.64 | 10.43 | 18.69 | 8.00 | 0.71 | 46.15 | 18.49 | 19.36 | 8.48 | 0.54 | 45.61 | 9.27 | 16.42 | 9.02 | 0.82 |
| | | | **5-HT→ Glu (+)** | 46.15 | 20.06 | 18.18 | 8.44 | 0.52 | 45.69 | 18.45 | 18.44 | 14.03 | 0.85 | 46.15 | 18.87 | 19.39 | 8.65 | 0.55 | 45.66 | 16.22 | 16.32 | 14.83 | 0.96 |
| b | **5-HT→Glu (-)** | | | 46.15 | 13.89 | 8.57 | 5.68 | 0.47 | 45.60 | 9.36 | 12.69 | 6.63 | 0.68 | 46.15 | 13.40 | 9.03 | 5.86 | 0.49 | 45.58 | 8.14 | 15.43 | 7.16 | 0.78 |
| | **5-HT→Glu (+)** | | | 46.15 | 13.08 | 8.64 | 5.31 | 4.60 | 45.54 | 9.13 | 12.80 | 6.35 | 6.71 | 46.15 | 12.68 | 9.11 | 5.48 | 4.86 | 45.51 | 8.07 | 15.54 | 6.90 | 7.75 |
| c | - | | | 46.15 | 12.16 | 8.71 | 4.95 | 0.00 | 45.60 | 8.18 | 12.96 | 5.71 | 0.00 | 46.15 | 11.74 | 9.18 | 5.11 | 0.00 | 45.57 | 7.12 | 15.74 | 6.19 | 0.00 |
| d | - | | | 46.15 | 13.36 | 1.28 | 4.71 | 0.00 | 45.60 | 8.88 | 0.27 | 5.67 | 0.00 | 46.15 | 12.85 | 1.19 | 4.88 | 0.00 | 45.57 | 7.72 | 0.63 | 6.16 | 0.00 |
| e | - | | | 46.15 | 12.40 | 2.12 | 4.31 | 0.00 | 45.58 | 8.40 | 3.95 | 5.28 | 0.00 | 46.15 | 11.97 | 2.24 | 4.47 | 0.00 | 45.55 | 7.33 | 4.61 | 5.76 | 0.00 |
| f | - | | | 46.15 | 1.03 | 0.16 | 0.32 | 0.00 | 45.52 | 0.57 | 0.26 | 0.35 | 0.00 | 46.15 | 0.94 | 0.17 | 0.34 | 0.00 | 45.49 | 0.46 | 0.30 | 0.38 | 0.00 |
| g | - | | | 46.15 | 0.00 | 7.18 | 0.00 | 0.00 | 45.52 | 0.00 | 8.64 | 0.00 | 0.00 | 46.15 | 0.00 | 7.28 | 0.00 | 0.00 | 45.49 | 0.00 | 9.30 | 0.00 | 0.00 |
| h | - | | | 46.15 | 13.36 | 2.31 | 4.71 | 0.00 | 45.58 | 8.90 | 4.24 | 5.68 | 0.00 | 46.15 | 12.85 | 2.45 | 4.88 | 0.00 | 45.56 | 7.73 | 4.93 | 6.17 | 0.00 |
| i | - | | | 46.15 | 0.00 | 0.00 | 0.00 | 0.00 | 45.49 | 0.00 | 0.00 | 0.00 | 0.00 | 46.15 | 0.00 | 0.00 | 0.00 | 0.00 | 45.46 | 0.00 | 0.00 | 0.00 | 0.00 |
| j | - | | | 46.15 | 0.00 | 0.00 | 0.00 | 0.00 | 45.55 | 0.00 | 0.00 | 0.00 | 0.00 | 46.15 | 0.00 | 0.00 | 0.00 | 0.00 | 45.52 | 0.00 | 0.00 | 0.00 | 0.00 |
| k | - | | | 46.15 | 0.00 | 0.00 | 0.00 | 0.00 | 45.52 | 0.00 | 0.00 | 0.00 | 0.00 | 46.15 | 0.00 | 0.00 | 0.00 | 0.00 | 45.49 | 0.00 | 0.00 | 0.00 | 0.00 |
| l | **5-HT→DA (-)** | | | 46.15 | 12.09 | 7.97 | 4.67 | 0.45 | 45.60 | 8.16 | 6.85 | 5.64 | 0.66 | 46.15 | 11.66 | 8.42 | 4.88 | 0.48 | 45.57 | 7.19 | 12.94 | 6.24 | 0.76 |
| | **5-HT→DA (+)** | | | 46.15 | 12.09 | 7.97 | 4.67 | 0.45 | 45.58 | 8.18 | 6.86 | 5.65 | 0.66 | 46.15 | 11.66 | 8.43 | 4.88 | 0.48 | 45.56 | 7.20 | 12.93 | 6.25 | 0.76 |

**Table 7. Percentage changes in neural population firing rates with $D_2$ receptor-mediated connection strengths changed by a factor of 40 (X=40).** Time-averaged percentage changes in neural population activities. Arc: Architecture type. Nomenclatures same as in Supplementary Information, Table 4, except GA, which is an abbreviation of GABA. Values in red: Percentage changes beyond the prescribed allowed ranges set by the inclusion criterion.

| Arc | | | | Type-I 5-HT neurons | | | | | | | | | | Type-II 5-HT neurons | | | | | | | | | |
|---|---|---|---|---|---|---|---|---|---|---|---|---|---|---|---|---|---|---|---|---|---|---|---|
| a | Connections/Type | | | **Punishment** | | | | | **Reward** | | | | | **Punishment** | | | | | **Reward** | | | | |
| | | | | **DA VTA** | **5HT DRN** | **GA DRN** | **GA VTA** | **Glu VTA** | **DA VTA** | **5HT DRN** | **GA DRN** | **GA VTA** | **Glu VTA** | **DA VTA** | **5HT DRN** | **GA DRN** | **GA VTA** | **Glu VTA** | **DA VTA** | **5HT DRN** | **GA DRN** | **GA VTA** | **Glu VTA** |
| | **DA→ GA-VTA (-)** | **5-HT→ DA (-)** | **5-HT→ Glu (-)** | 46.15 | 37.21 | 7.93 | 15.93 | 0.69 | 46.15 | 30.28 | 23.34 | 23.77 | 1.12 | 46.15 | 35.33 | 8.35 | 16.24 | 0.72 | 46.15 | 26.75 | 18.91 | 23.62 | 1.19 |
| | | | **5-HT→ Glu (+)** | 46.15 | 37.67 | 7.87 | 16.21 | 0.70 | 46.14 | 20.93 | 23.51 | 15.63 | 0.93 | 46.15 | 35.80 | 8.29 | 16.52 | 0.73 | 46.14 | 18.59 | 19.01 | 16.55 | 1.03 |
| | **5-HT→ GA-VTA (-)** | **5-HT→ DA (+)** | **5-HT→ Glu (-)** | 46.15 | 37.21 | 7.93 | 15.93 | 0.69 | 46.15 | 20.53 | 23.52 | 15.28 | 0.92 | 46.15 | 35.33 | 8.35 | 16.24 | 0.72 | 46.14 | 18.19 | 19.01 | 16.21 | 1.02 |
| | | | **5-HT→ Glu (+)** | 46.15 | 37.67 | 7.87 | 16.21 | 0.70 | 46.15 | 34.34 | 23.65 | 27.30 | 1.20 | 46.15 | 35.80 | 8.29 | 16.52 | 0.73 | 46.15 | 30.49 | 19.10 | 26.86 | 1.26 |
| | **DA→ GA-VTA (+)** | **5-HT→ DA (-)** | **5-HT→ Glu (-)** | 46.15 | 36.38 | 22.50 | 17.96 | 0.69 | 46.15 | 30.16 | 24.75 | 25.61 | 1.11 | 46.15 | 34.52 | 23.79 | 18.27 | 0.71 | 46.15 | 26.65 | 19.78 | 25.51 | 1.19 |
| | | | **5-HT→ Glu (+)** | 46.15 | 36.83 | 22.55 | 18.24 | 0.69 | 46.15 | 20.56 | 24.68 | 17.23 | 0.92 | 46.15 | 34.98 | 23.84 | 18.55 | 0.72 | 46.14 | 18.26 | 19.74 | 18.21 | 1.03 |
| | **5-HT→ GA-VTA (+)** | **5-HT→ DA (+)** | **5-HT→ Glu (-)** | 46.15 | 36.38 | 22.50 | 17.96 | 0.69 | 46.15 | 20.16 | 24.67 | 16.89 | 0.91 | 46.15 | 34.52 | 23.79 | 18.27 | 0.71 | 46.15 | 17.86 | 19.74 | 17.88 | 1.02 |
| | | | **5-HT→ Glu (+)** | 46.15 | 36.83 | 22.55 | 18.24 | 0.69 | 46.15 | 34.14 | 24.40 | 29.08 | 1.19 | 46.15 | 34.98 | 23.84 | 18.55 | 0.72 | 46.15 | 30.31 | 19.56 | 28.69 | 1.26 |
| b | **5-HT→Glu (-)** | | | 46.15 | 23.69 | 10.39 | 10.62 | 0.55 | 46.13 | 15.30 | 14.22 | 11.77 | 0.79 | 46.15 | 22.82 | 10.86 | 10.90 | 0.58 | 46.12 | 13.36 | 17.06 | 12.39 | 0.89 |
| | **5-HT→Glu (+)** | | | 46.15 | 23.08 | 10.47 | 10.26 | 5.42 | 46.11 | 15.20 | 14.29 | 11.57 | 7.84 | 46.15 | 22.30 | 10.94 | 10.54 | 5.70 | 46.10 | 13.41 | 17.13 | 12.20 | 8.88 |
| c | - | | | 46.15 | 22.01 | 10.56 | 9.76 | 0.00 | 46.13 | 14.15 | 14.52 | 10.76 | 0.00 | 46.15 | 21.21 | 11.04 | 10.02 | 0.00 | 46.12 | 12.36 | 17.38 | 11.37 | 0.00 |
| d | - | | | 46.15 | 23.23 | 1.81 | 9.09 | 0.00 | 46.13 | 14.87 | 0.60 | 10.40 | 0.00 | 46.15 | 22.34 | 1.67 | 9.36 | 0.00 | 46.12 | 12.97 | 1.04 | 10.99 | 0.00 |
| e | - | | | 46.15 | 22.32 | 4.24 | 8.63 | 0.00 | 46.12 | 14.42 | 7.45 | 9.99 | 0.00 | 46.15 | 21.52 | 4.46 | 8.90 | 0.00 | 46.11 | 12.62 | 8.40 | 10.59 | 0.00 |
| f | - | | | 46.15 | 1.03 | 0.16 | 0.32 | 0.00 | 46.08 | 0.57 | 0.26 | 0.35 | 0.00 | 46.15 | 0.94 | 0.17 | 0.34 | 0.00 | 46.07 | 0.46 | 0.30 | 0.38 | 0.00 |
| g | - | | | 46.15 | 0.00 | 12.56 | 0.00 | 0.00 | 46.08 | 0.00 | 14.50 | 0.00 | 0.00 | 46.15 | 0.00 | 12.74 | 0.00 | 0.00 | 46.07 | 0.00 | 15.55 | 0.00 | 0.00 |
| h | - | | | 46.15 | 23.23 | 4.47 | 9.09 | 0.00 | 46.12 | 14.87 | 7.76 | 10.41 | 0.00 | 46.15 | 22.34 | 4.70 | 9.36 | 0.00 | 46.11 | 12.98 | 8.72 | 11.00 | 0.00 |
| i | - | | | 46.15 | 0.00 | 0.00 | 0.00 | 0.00 | 46.07 | 0.00 | 0.00 | 0.00 | 0.00 | 46.15 | 0.00 | 0.00 | 0.00 | 0.00 | 46.05 | 0.00 | 0.00 | 0.00 | 0.00 |
| j | - | | | 46.15 | 0.00 | 0.00 | 0.00 | 0.00 | 46.09 | 0.00 | 0.00 | 0.00 | 0.00 | 46.15 | 0.00 | 0.00 | 0.00 | 0.00 | 46.08 | 0.00 | 0.00 | 0.00 | 0.00 |
| k | - | | | 46.15 | 0.00 | 0.00 | 0.00 | 0.00 | 46.08 | 0.00 | 0.00 | 0.00 | 0.00 | 46.15 | 0.00 | 0.00 | 0.00 | 0.00 | 46.07 | 0.00 | 0.00 | 0.00 | 0.00 |
| l | **5-HT→DA (-)** | | | 46.15 | 21.92 | 9.76 | 9.00 | 0.53 | 46.13 | 14.12 | 8.66 | 10.34 | 0.76 | 46.15 | 21.12 | 10.21 | 9.32 | 0.56 | 46.12 | 12.42 | 13.39 | 11.06 | 0.87 |
| | **5-HT→DA (+)** | | | 46.15 | 21.92 | 9.76 | 9.00 | 0.53 | 46.12 | 14.13 | 8.66 | 10.34 | 0.76 | 46.15 | 21.12 | 10.21 | 9.32 | 0.56 | 46.11 | 12.42 | 13.39 | 11.06 | 0.87 |

**Table 8. Percentage changes in neural population firing rates with $D_2$ receptor-mediated connection strengths changed by a factor of 70 (X=70).** Time-averaged percentage changes in neural population activities. Arc: Architecture type. Nomenclatures same as in Supplementary Information, Table 4, except GA, which is an abbreviation of GABA. Values in red: Percentage changes beyond the prescribed allowed ranges set by the inclusion criterion.

| Arc | Connections/Type | | | Type-I 5-HT neurons | | | | | | | | | | Type-II 5-HT neurons | | | | | | | | | |
|---|---|---|---|---|---|---|---|---|---|---|---|---|---|---|---|---|---|---|---|---|---|---|---|
| | | | | Punishment | | | | | Reward | | | | | Punishment | | | | | Reward | | | | |
| | | | | DA VTA | 5HT DRN | GA DRN | GA VTA | Glu VTA | DA VTA | 5HT DRN | GA DRN | GA VTA | Glu VTA | DA VTA | 5HT DRN | GA DRN | GA VTA | Glu VTA | DA VTA | 5HT DRN | GA DRN | GA VTA | Glu VTA |
| a | DA→ GA-VTA (-) | 5-HT→ DA (-) | 5-HT→ Glu (-) | 46.15 | 53.13 | 8.20 | 26.88 | 0.91 | 46.15 | 41.52 | 29.35 | 32.48 | 1.33 | 46.15 | 50.73 | 8.65 | 27.08 | 0.94 | 46.15 | 36.94 | 23.08 | 31.05 | 1.37 |
| | | | 5-HT→ Glu (+) | 46.15 | 53.71 | 8.11 | 27.31 | 0.92 | 46.15 | 31.00 | 29.51 | 24.62 | 1.14 | 46.15 | 51.31 | 8.56 | 27.50 | 0.94 | 46.15 | 27.69 | 23.19 | 24.71 | 1.23 |
| | 5-HT→ GA-VTA (-) | 5-HT→ DA (+) | 5-HT→ Glu (-) | 46.15 | 53.13 | 8.20 | 26.88 | 0.91 | 46.15 | 30.52 | 29.52 | 24.21 | 1.14 | 46.15 | 50.73 | 8.65 | 27.08 | 0.94 | 46.15 | 27.21 | 23.20 | 24.35 | 1.22 |
| | | | 5-HT→ Glu (+) | 46.15 | 53.71 | 8.11 | 27.31 | 0.92 | 46.15 | 50.86 | 29.70 | 37.51 | 1.44 | 46.15 | 51.31 | 8.56 | 27.50 | 0.94 | 46.15 | 45.75 | 23.39 | 35.36 | 1.46 |
| | DA→ GA-VTA (+) | 5-HT→ DA (-) | 5-HT→ Glu (-) | 46.15 | 52.04 | 27.22 | 29.69 | 0.89 | 46.15 | 41.36 | 31.02 | 35.15 | 1.33 | 46.15 | 49.65 | 28.56 | 29.91 | 0.92 | 46.15 | 36.79 | 24.52 | 33.79 | 1.37 |
| | | | 5-HT→ Glu (+) | 46.15 | 52.60 | 27.29 | 30.11 | 0.90 | 46.15 | 30.49 | 30.94 | 26.94 | 1.13 | 46.15 | 50.22 | 28.63 | 30.31 | 0.93 | 46.15 | 27.22 | 24.46 | 27.13 | 1.22 |
| | 5-HT→ GA-VTA (+) | 5-HT→ DA (+) | 5-HT→ Glu (-) | 46.15 | 52.04 | 27.22 | 29.69 | 0.89 | 46.15 | 30.01 | 30.94 | 26.52 | 1.12 | 46.15 | 49.65 | 28.56 | 29.91 | 0.92 | 46.15 | 26.74 | 24.46 | 26.77 | 1.21 |
| | | | 5-HT→ Glu (+) | 46.15 | 52.60 | 27.29 | 30.11 | 0.90 | 46.15 | 50.57 | 30.61 | 40.17 | 1.44 | 46.15 | 50.22 | 28.63 | 30.31 | 0.93 | 46.15 | 45.48 | 24.15 | 38.10 | 1.46 |
| b | 5-HT→Glu (-) | | | 46.15 | 33.35 | 12.06 | 16.38 | 0.65 | 46.15 | 21.43 | 15.72 | 17.51 | 0.91 | 46.15 | 32.12 | 12.54 | 16.72 | 0.68 | 46.15 | 18.72 | 18.76 | 17.97 | 1.01 |
| | 5-HT→Glu (+) | | | 46.15 | 32.96 | 12.12 | 16.06 | 6.40 | 46.15 | 21.50 | 15.75 | 17.42 | 9.11 | 46.15 | 31.81 | 12.60 | 16.40 | 6.69 | 46.15 | 18.91 | 18.78 | 17.90 | 10.09 |
| c | - | | | 46.15 | 31.71 | 12.26 | 15.38 | 0.00 | 46.15 | 20.32 | 16.04 | 16.44 | 0.00 | 46.15 | 30.55 | 12.74 | 15.71 | 0.00 | 46.15 | 17.76 | 19.08 | 16.95 | 0.00 |
| d | - | | | 46.15 | 32.95 | 1.95 | 14.29 | 0.00 | 46.15 | 21.03 | 1.41 | 15.73 | 0.00 | 46.15 | 31.70 | 1.77 | 14.62 | 0.00 | 46.15 | 18.37 | 1.71 | 16.19 | 0.00 |
| e | - | | | 46.15 | 32.11 | 6.77 | 13.78 | 0.00 | 46.15 | 20.64 | 11.42 | 15.32 | 0.00 | 46.15 | 30.94 | 7.07 | 14.11 | 0.00 | 46.15 | 18.06 | 12.47 | 15.79 | 0.00 |
| f | - | | | 46.15 | 1.03 | 0.16 | 0.32 | 0.00 | 46.15 | 0.57 | 0.26 | 0.35 | 0.00 | 46.15 | 0.94 | 0.17 | 0.34 | 0.00 | 46.15 | 0.46 | 0.30 | 0.38 | 0.00 |
| g | - | | | 46.15 | 0.00 | 17.95 | 0.00 | 0.00 | 46.15 | 0.00 | 20.63 | 0.00 | 0.00 | 46.15 | 0.00 | 18.20 | 0.00 | 0.00 | 46.15 | 0.00 | 22.12 | 0.00 | 0.00 |
| h | - | | | 46.15 | 32.95 | 7.02 | 14.29 | 0.00 | 46.15 | 21.03 | 11.73 | 15.73 | 0.00 | 46.15 | 31.70 | 7.33 | 14.62 | 0.00 | 46.15 | 18.37 | 12.77 | 16.19 | 0.00 |
| i | - | | | 46.15 | 0.00 | 0.00 | 0.00 | 0.00 | 46.15 | 0.00 | 0.00 | 0.00 | 0.00 | 46.15 | 0.00 | 0.00 | 0.00 | 0.00 | 46.15 | 0.00 | 0.00 | 0.00 | 0.00 |
| j | - | | | 46.15 | 0.00 | 0.00 | 0.00 | 0.00 | 46.15 | 0.00 | 0.00 | 0.00 | 0.00 | 46.15 | 0.00 | 0.00 | 0.00 | 0.00 | 46.15 | 0.00 | 0.00 | 0.00 | 0.00 |
| k | - | | | 46.15 | 0.00 | 0.00 | 0.00 | 0.00 | 46.15 | 0.00 | 0.00 | 0.00 | 0.00 | 46.15 | 0.00 | 0.00 | 0.00 | 0.00 | 46.15 | 0.00 | 0.00 | 0.00 | 0.00 |
| l | 5-HT→DA (-) | | | 46.15 | 31.61 | 11.38 | 14.13 | 0.63 | 46.15 | 20.27 | 10.45 | 15.64 | 0.89 | 46.15 | 30.44 | 11.84 | 14.52 | 0.66 | 46.15 | 17.79 | 13.82 | 16.26 | 0.99 |
| | 5-HT→DA (+) | | | 46.15 | 31.61 | 11.38 | 14.13 | 0.63 | 46.15 | 20.27 | 10.45 | 15.64 | 0.89 | 46.15 | 30.44 | 11.84 | 14.52 | 0.66 | 46.15 | 17.79 | 13.82 | 16.26 | 0.99 |

**Table 9. Percentage changes in neural population firing rates with $D_2$ receptor-mediated connection strengths changed by a factor of 100 (X=100).** Time-averaged percentage changes in neural population activities. Arc: Architecture type. Nomenclatures same as in Supplementary Information, Table 4, except GA, which is an abbreviation of GABA. Values in red: Percentage changes beyond the prescribed allowed ranges set by the inclusion criterion.

### Supplementary Method

#### Mathematical derivation for steady states (fixed points) in neural circuit models

For each degenerate circuit model, local stability analysis was used to find the stability of a system of dynamical equations that describe DRN-VTA circuit dynamics. The local stability of the network can be determined by first determining the steady states (also called fixed points) and then identifying whether each of these fixed points, if they exist, are stable (Strogatz, 2018).

The computational models to be investigated were based on our previous mean-field, neural population-based modelling framework for neuromodulator circuits (Joshi et al., 2017), in which the averaged concentration releases of neuromodulators ([5-HT] and [DA]) were monotonic functions of the averaged firing rate of (5-HT and DA) neuronal populations via some neuromodulator induced slow currents.

Each network's steady state (or equilibrium/fixed point) can be obtained by setting the rate of change for all the above dynamical equations (Eqs. (25-30)) to zero, i.e. $\frac{dI_{auto,5-HT}}{dt} = \frac{dI_{auto,DA}}{dt} = \frac{dI_{DA,5-HT}}{dt} = \frac{dI_{5-HT,DA}}{dt} = \frac{d[5-HT]}{dt} = \frac{d[DA]}{dt} = 0$, and then solving them algebraically. The solution of these equations will give the steady-state value for each model. Specifically, the currents (dynamical variables) from Eqs. (25-30) in the main text (e.g. $I_{auto,5-HT} = \frac{80}{1+e^{-10([5-HT]-0.1)}}$) were substituted into the input-output functions described by Eqs. (20-24) in the main text. Note that the different model architectures (Fig. 4 and Supplementary Information, Table 4) can be determined from the specific sets of values of $W$'s and $J$'s, and the biased currents. Henceforth, only the generic solution is provided. The steady-state values for the model can be re-written using Eqs. (25-30), such that

$$I_{auto,5-HT} = \frac{80}{1+x} \quad (34)$$

$$I_{auto,DA} = \frac{80}{1+y} \quad (35)$$

$$I_{DA,5-HT} = \frac{0.03}{1+x^2} \quad (36)$$

$$I_{5-HT,DA} = \frac{0.03}{1+ay^2} \quad (37)$$

$$F_{DA} = \frac{40[DA]}{0.15+[DA]} \quad (38)$$

$$F_{5-HT} = \frac{16.25[5HT]}{0.17+[5HT]} \quad (39)$$

where $x = e^{-10([5-HT]-0.1)}$,$y = e^{-10([DA]-0.1)}$ and with $a = e^4$. Hence, at the steady state, considering all-to-all connectivity (the most general case), and using the explicit parameter values and Eqs. (34-39) above, the afferent input currents from Eqs. (9-13) in the main text can be written as

$$I_{DA} = -J_{auto,DA}\frac{80}{1+y} \pm W_{d5}\frac{0.03}{1+x^2} + J_{d5e}F_{Glu} - J_{di}F_{GABA-VTA} - J_{d5i}F_{GABA-DRN} + 210 + I_{DA,ext} \quad (40)$$

$$I_{5-HT} = -J_{auto,5-HT}\frac{80}{1+x} + W_{5d}\frac{0.03}{1+ay^2} + J_{55e}F_{Glu} - J_{55i}F_{GABA-DRN} - J_{5i}F_{GABA-VTA} + 99.87 + I_{5-HT,ext} \quad (41)$$

$$I_{Glu} = J_{self,Glu}F_{Glu} \pm W_{5e5}\frac{0.03}{1+x^2} - W_{5edi}F_{GABA-VTA} + W_{5e2}\frac{0.03}{1+ay^2} + 100 + I_{Glu,ext} \quad (42)$$

$$I_{GABA-DRN} = -J_{self,GABA-DRN}F_{GABA-DRN} \pm W_{5i5}\frac{0.03}{1+x^2} + W_{5id}\frac{0.03}{1+ay^2} + W_{5ie} * F_{Glu} - W_{5ii}I_{GABA-DRN,GABA-VTA} + 450 + I_{GABA-DRN,ext} \quad (43)$$

$$I_{GABA-VTA} = -J_{self,GABA-VTA}F_{GABA-VTA} + W_{i5}\frac{0.03}{1+x^2} \pm W_{id}\frac{0.03}{1+ay^2} + 200 + I_{GABA-VTA,ext} \quad (44)$$

Now, using only the linear parts of the above threshold-linear functions (which were validated post-hoc), the firing rates in Eqs. (20-24) can be written, after some algebraic manipulations, as

$$F_{DA} = g_{DA}\left(-J_{auto,DA}\frac{80}{1+y} \pm W_{d5}\frac{0.03}{1+x^2} + J_{d5e}F_{Glu} - J_{di}F_{GABA-VTA} - J_{d5i}F_{GABA-DRN} + 210 + I_{DA,ext}\right) \quad (45)$$

$$F_{5-HT} = g_{5-HT}\left(-J_{auto,5-HT}\frac{80}{1+x} + W_{5d}\frac{0.03}{1+ay^2} + J_{55e}F_{Glu} - J_{55i}F_{GABA-DRN} - J_{5i}F_{GABA-VTA} + 99.87 + I_{5-HT,ext}\right) \quad (46)$$

$$F_{Glu} = g_{Glu}[J_{self,Glu}F_{Glu} \pm W_{5e5}I_{Glu,5-HT} + 100 + I_{Glu,ext}]_+ \quad (47)$$

$$F_{GABA-DRN} = g_{GABA-DRN}[-J_{self,GABA-DRN}F_{GABA-DRN} \pm W_{5i5}I_{GABA-DRN,5-HT} + W_{5id}I_{GABA-DRN,DA} - W_{5ii}I_{GABA-DRN,GABA-VTA} + 450 + I_{GABA-DRN,ext}]_+ \quad (48)$$

$$F_{GABA-VTA} = g_{GABA-VTA}[-J_{self,GABA-VTA}\, F_{GABA-VTA} + W_{i5}\, I_{GABA-VTA,5-HT} \pm W_{id}\, I_{GABA-VTA,DA} + 200 + I_{GABA-VTA,ext}]_+ \quad (49)$$

Eqs. (45-49) can then be rewritten in matrix form, as shown in Eq. (31-32) in the main text.
